# EDA2R/NIK signaling promotes skeletal muscle atrophy linked to cancer cachexia

**DOI:** 10.1101/2023.01.23.525138

**Authors:** Sevval Nur Bilgic, Aylin Domaniku, Batu Toledo, Samet Agca, Bahar Z. C. Weber, Dilsad H. Arabaci, Zeynep Ozornek, Pascale Lause, Jean-Paul Thissen, Audrey Loumaye, Serkan Kir

**Affiliations:** Department of Molecular Biology and Genetics, Koc University, Istanbul 34450, Turkey; Department of Endocrinology and Nutrition, Cliniques Universitaires Saint-Luc, 1200 Brussels Belgium; Pole of Endocrinology, Diabetology and Nutrition, Institute of Experimental and Clinical Research, Université Catholique de Louvain, 1200 Brussels, Belgium

## Abstract

Skeletal muscle atrophy is a hallmark of the cachexia syndrome that is associated with poor survival and reduced quality of life in cancer patients^1^. Muscle atrophy involves excessive protein catabolism and loss of muscle mass and strength^2^. An effective therapy against muscle wasting is lacking as mechanisms driving the atrophy process remain incompletely understood. Our gene expression analysis in muscle tissues revealed upregulation of Ectodysplasin A2 Receptor (EDA2R) in tumor-bearing mice and cachectic cancer patients. Here we show that activation of EDA2R signaling promotes skeletal muscle atrophy. Stimulation of primary myotubes with EDA2R ligand, EDA-A2, triggered pronounced cellular atrophy via inducing the expression of muscle atrophy-related genes *Atrogin1* and *MuRF1*. EDA-A2-driven myotube atrophy involved activation of the noncanonical NFκB pathway and depended on NIK kinase activity. While EDA-A2 overexpression induced muscle wasting in mice, the deletion of EDA2R or muscle NIK protected tumor-bearing mice from the loss of muscle mass and function. Tumor-induced Oncostatin M upregulated muscle EDA2R expression and muscle-specific Oncostatin M Receptor (OSMR) knockout mice were resistant to tumor-driven muscle wasting. Our results demonstrate that EDA2R/NIK signaling mediates cancer-associated muscle atrophy in an OSM/OSMR-dependent manner. Thus, therapeutic targeting of these pathways may be beneficial in preventing muscle loss.

---

Skeletal muscle atrophy is characterized by excessive protein catabolism leading to loss of muscle mass and strength^2^. Muscle loss is associated with aging (i.e., sarcopenia), muscular dystrophies and the cachexia syndrome that is linked to chronic diseases such as cancer and kidney failure. Cachexia involves progressive muscle wasting that is often accompanied by the loss of adipose tissue^3^. Cachexia is highly prevalent in patients with lung, gastric, pancreatic or colorectal cancers where it leads to dramatic weight loss and poor quality of life. Cancer patients experiencing cachexia exhibit frailty, reduced response to treatment and poor survival^1^. Without an effective therapy to reverse muscle wasting, cachexia remains a major problem for the patients^4^. A better understanding of tumor-driven mechanisms promoting muscle atrophy is urgently needed to design new therapeutics.

Ectodysplasin A (EDA) is a Tumor Necrosis Factor (TNF) family member involved in ectodermal development. Mutations in the *EDA* gene have been associated with X-linked hypohidrotic ectodermal dysplasia, a congenital disease characterized by abnormalities in the development of skin, hair, nails, teeth, and sweat glands^5^. Alternative splicing generates numerous *EDA* transcripts, including well-known isoforms EDA-A1 and EDA-A2 which differ only by two amino acids missing in EDA-A2. While EDA-A1 binds to the EDAR receptor, EDA-A2 exclusively interacts with EDA2R^6^. These ligands are produced as transmembrane proteins but they are also enzymatically cleaved and secreted. Defects in *EDA* and *EDAR* genes are linked to ectodermal dysplasia^7^. However, the deletion of EDA2R does not affect the development of ectodermal tissues and the physiological roles of the EDA-A2/EDA2R pathway remain largely elusive^8^.

We investigated skeletal muscle wasting driven by cachexia-inducing Lewis Lung Carcinoma (LLC) and B16 melanoma tumors in the syngeneic C57BL/6 mice^9,10^. Our gene expression analysis revealed significant upregulation of *Eda2r* mRNA in skeletal muscles of tumor-bearing mice (Fig. 1a). In fact, by comparing gene expression in different tissues of mice, we identified high levels of *Eda-a1* and *Eda-a2* mRNA in skeletal muscle (Extended Data Fig. 1a,b). While *Edar* expression was the highest in the skin, *Eda2r* mRNA levels were markedly enriched in skeletal muscle (Extended Data Fig. 1c,d), arguing a potential role for EDA2R signaling in muscle pathophysiology. Testing gene expression in muscle biopsies, we detected elevated *EDA2R* mRNA in a subset of lung and colorectal cancer patients experiencing cachexia (Fig. 1b). Furthermore, our analysis of gene expression datasets indicated significantly increased *EDA2R* transcript levels in muscle biopsies of cachectic patients with pancreatic ductal adenocarcinoma (PDAC) compared to non-cachectic PDAC patients and non-cancer subjects (Fig. 1c). Similarly, muscle EDA2R levels were elevated in cachectic patients with upper gastrointestinal cancers (UGIC) compared to healthy controls. Upon the resection of tumors, there was a trend for reduced *EDA2R* expression in these patients (Fig. 1d and Extended Data Fig. 2a). In addition, we also detected significantly elevated *EDA2R* transcript in muscle biopsies of Duchenne muscular dystrophy (DMD) and Facioscapulohumeral muscular dystrophy (FSHD) patients who suffer from reduced muscle mass and function (Extended Data Fig. 2b-d). Intrigued by these observations, we asked whether the EDA-A2/EDA2R pathway is involved in muscle loss.

**Figure 1.**
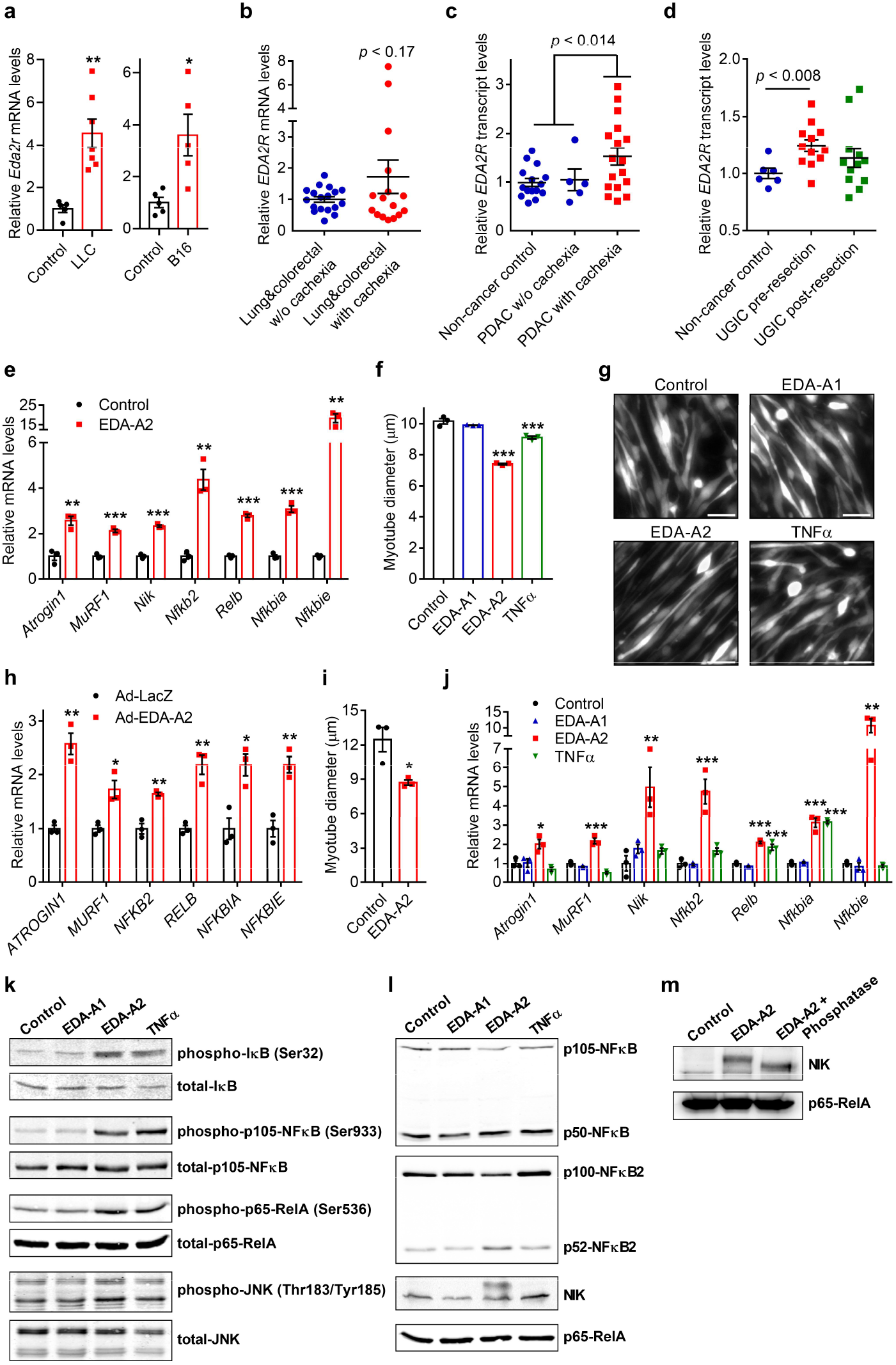
EDA-A2 promotes atrophy and activates NFκB signaling in myotubes. **a,** *Eda2r* mRNA levels were tested by RT-qPCR in gastrocnemius muscles of mice bearing LLC tumors (for 16 days) (control n = 5, LLC n = 7) or B16 tumors (for 14 days) (n = 5). **b**, *EDA2R* mRNA levels were determined by RT-qPCR in quadriceps muscle biopsies of lung and colorectal cancer patients with or without cachexia (w/o cachexia n = 16, with cachexia n = 18). **c**, *EDA2R* transcript levels were analyzed in rectus abdominis muscle biopsies from non-cancer controls (n = 15), and PDAC patients with cachexia (n = 17) and without cachexia (n = 5) (GSE130563). **d**, *EDA2R* transcript levels were analyzed in quadriceps muscle biopsies collected from non-cancer subjects (n = 6) and cachectic patients with upper gastrointestinal cancer (n = 12) before and after tumor resection (GSE34111). **e**, Fully differentiated mouse primary myotubes were treated with recombinant EDA-A2 (250 ng/ml) for 24 hr and gene expression was determined by RT-qPCR (n = 3). **f**,**g**, Mouse primary myotubes were treated with recombinant EDA-A1, EDA-A2 or TNFα proteins (250 ng/ml each) for 48 hr. Cells also transduced with a GFP adenovirus were visualized under the fluorescence microscope. Average myotube diameter was measured (n = 3) (**f**). The scale bar is 50 μm (**g**). **h**,**i**, Human Skeletal Muscle Myoblasts (HSMM) were differentiated into myotubes and treated with an adenovirus expressing EDA-A2 (**h**) or recombinant EDA-A2 protein (250 ng/ml) (**i**) for 48 hr. Gene expression was determined by RT-qPCR (n = 3) (**h**) and myotube diameter was measured (n = 3) (**i**). **j-m**, Mouse primary myotubes were treated with EDA-A1, EDA-A2 or TNFα (250 ng/ml each) for 24 hr (**j**,**l**,**m**) or 10 min (**k**). Changes in gene expression were determined by RT-qPCR (n = 3) (**j**). Cell lysates were investigated by western blotting (**k**,**l**). The lysate of EDA-A2-treated myotubes was also treated with alkaline phosphatase (**m**). The values are mean ± SEM. Statistical analysis was conducted using the two-tailed t-test (**a**,**b**,**c**,**d**,**e**,**h**,**i**) and one-way ANOVA (**f**,**j**). \**p* < 0.05, \*\**p* < 0.01, \*\*\**p* < 0.001, compared with the Control group.

First, we studied the outcome of the activation of this pathway in muscle cells. For this purpose, we isolated primary myoblasts from C57BL/6 mice and differentiated them into fully mature myotube cells. Treatment of primary myotubes with recombinant EDA-A2 protein stimulated mRNA levels of muscle atrophy-related genes; *Atrogin1* (*Fbxo32*) and *MuRF1* (*Trim63*) (Fig. 1e). These genes encode E3 ubiquitin ligase enzymes that are well-recognized inducers of muscle protein breakdown^2^. EDA-A2 treatment promoted cellular atrophy in primary myotubes as evidenced by a reduction in the diameter of these cells (Fig. 1f,g). While primary myotubes did not respond to EDA-A1 treatment, a marginal effect on cellular atrophy was induced by TNFα (Fig. 1f,g). Overexpression of EDA-A2 and not EDA-A1 in primary myotubes promoted mRNA levels of *Atrogin1* and *MuRF1*, and induced cellular atrophy (Extended Data Fig. 3a-c). In addition, the overexpression of EDA-A2 or the administration of recombinant EDA-A2 also induced *ATROGIN1* and *MURF1* expression in human myotubes and led to a reduction in myotube diameter (Fig. 1h,i and Extended Data Fig. 3d-f). We further documented the atrophic effects of EDA-A2 in mouse primary myotubes by measuring myosin heavy chain (MyHC) protein levels. Immunofluorescently labeled MyHC signal dropped significantly post EDA-A2 administration (Extended Data Fig 3g,h). MyHC downregulation was also detected by western blotting. Notably, proteasomal inhibition by MG132 reversed this effect, arguing that EDA-A2 promoted MyHC loss by enhancing proteasomal degradation (Extended Data Fig 3i).

Previous studies indicated the activation of NFκB signaling by EDA-A2/EDA2R^11,12^. Therefore, we further tested changes in mRNA and protein levels of NFκB factors and their IκB inhibitors. NFκB transcription factors are normally sequestered in the cytoplasm by inhibitory IκB proteins. The canonical NFκB signaling involves IKKβ-dependent phosphorylation of IκBs and their degradation while the noncanonical (alternative) NFκB activation depends on NFκB-inducing kinase (NIK) which promotes the processing of p100-NFκB2 into the active p52-NFκB2 form resulting in nuclear translocation of the p52-NFκB2/RelB complex^13^. Notably, EDA-A2 administration or overexpression increased mRNA levels of *Nik (Map3k14), Nfkb2, Relb, Nfkbia* and *Nfkbie* in mouse primary myotubes (Fig. 1e and Extended Data Fig. 3a and 4a,b). A similar effect was also detected in human myotubes (Fig. 1h and Extended Data Fig. 3e). Compared to TNFα, EDA-A2 treatment in mouse primary myotubes promoted a more pronounced effect on the expression of the atrophy genes and the NFκB signaling elements while EDA-A1 failed to stimulate these changes (Fig. 1j). Our results demonstrate that EDA-A2 induces the atrophy of myotubes and the transcription of noncanonical NFκB signaling components along with high levels of IκBs.

Next, we examined the activation of the canonical NFκB pathway by determining the phosphorylation of IκB, p105-NFκB and p65-RelA. Treatment of mouse primary myotubes with either EDA-A2 or TNFα induced the phosphorylation of these proteins acutely (Fig. 1k). We also detected an EDA-A2-induced increase in the phosphorylation of JNK, which was previously implicated in EDA2R signaling^11,14^ (Fig. 1k). A stable change in the processing of p105-NFκB into p50-NFκB was not detected after 24 hours of treatment (Fig. 1l). Interestingly, prolonged treatment of primary myotubes with recombinant EDA-A2 promoted alternative activation of NFκB signaling as evidenced by increased processing of p100-NFκB2 into p52-NFκB2, a process driven by NIK^13^. In fact, EDA-A2 treatment elevated NIK protein levels and particularly induced an electrophoretic mobility shift of this protein (Fig. 1l,m). This shift is in part due to the phosphorylation of NIK since it can be partially suppressed by alkaline phosphatase treatment (Fig. 1m). Other types of post-translational modifications likely contribute to this behavior. Wild-type mouse NIK protein overexpressed in primary myotubes also exhibited the mobility shift, unlike kinase-dead and autophosphorylation-deficient NIK mutants (Extended Data Fig. 4c). Notably, EDA-A2 treatment increased protein levels of the mutants without inducing a shift. Therefore, EDA-A2-induced mobility shift likely requires intact NIK kinase activity and depends on the autophosphorylation of the protein (Extended Data Fig. 4c). This shift was not detected for human NIK protein overexpressed in mouse primary myotubes (Extended Data Fig. 4d). In addition, overexpression of EDA-A2 in primary myotubes also activated the alternative NFκB signaling via NIK accumulation (Extended Data Fig. 4e). Our results indicate that EDA-A2 is a potent inducer of atrophy in myotubes where it leads to transient and stable activation of the canonical and noncanonical NFκB pathways, respectively.

To distinguish the relative contribution of NFκB pathways to EDA-A2-driven atrophy, we treated primary myotubes with the selective IKKβ inhibitor TPCA-1 and the proteasome inhibitor MG132. We found that EDA-A2-induced phosphorylation of IκB, p105-NFκB and p65-RelA was blocked by TPCA-1 treatment. However, inhibition of the canonical NFκB pathway did not suppress EDA-A2’s effects on gene expression (Fig. 2a,b). We tested additional IκB phosphorylation inhibitors such as BAY 11-7082 and BOT-64, which also failed to block EDA-A2-induced transcriptional changes (Extended Data Fig. 5a). IκB phosphorylation inhibitors also did not alter p100-NFκB2 processing driven by EDA-A2 (Fig. 2c and Extended Data Fig. 5b). On the other hand, proteasomal inhibition by MG132 was able to prevent p100-NFκB2 processing and the transcriptional effects elicited by EDA-A2 treatment in primary myotubes (Fig. 2a,c), consistent with an implication of the alternative NFκB activation at the downstream of EDA-A2 signaling.

**Figure 2.**
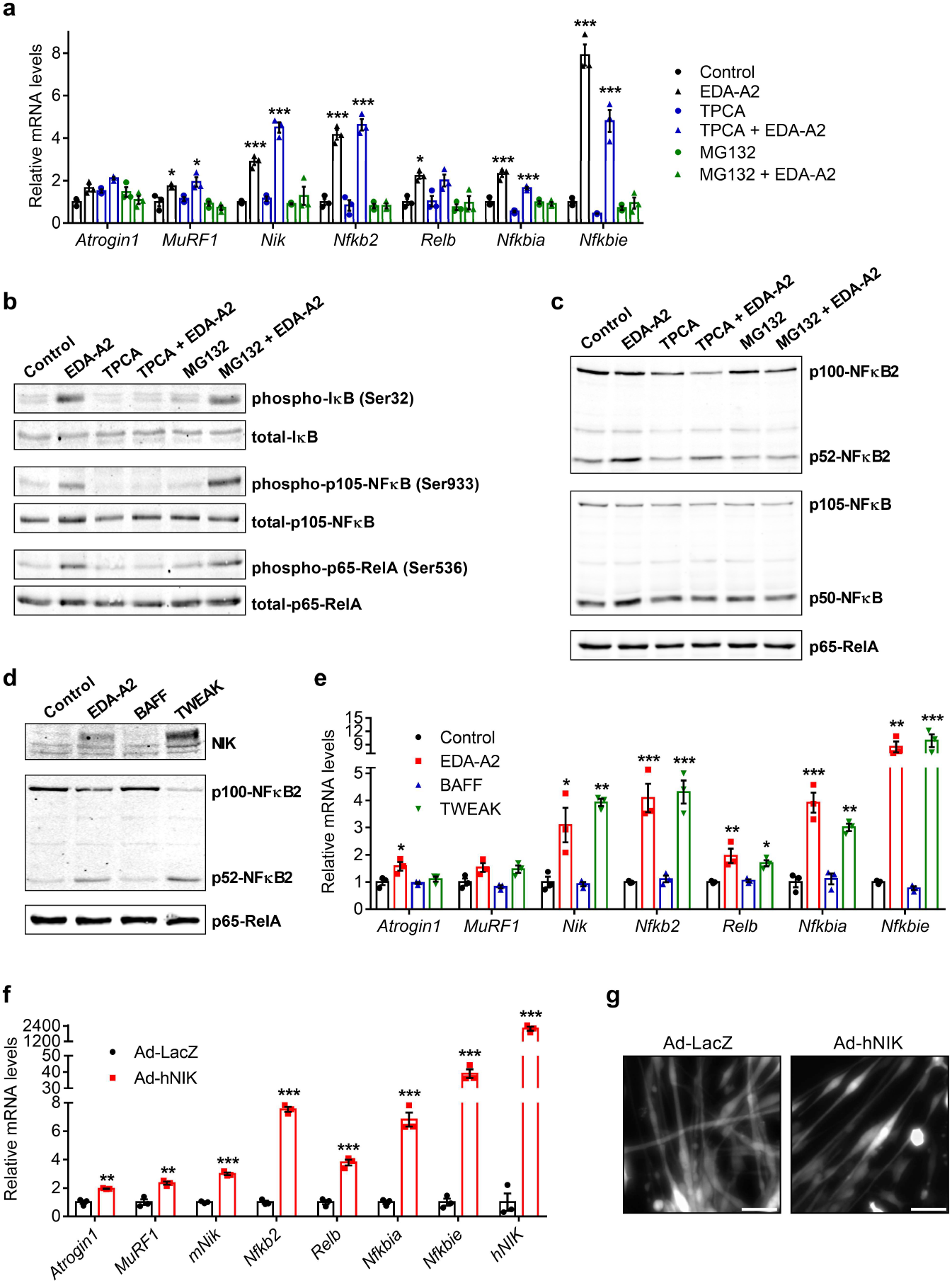
Activation of the noncanonical NFκB signaling by EDA-A2 or NIK kinase induces atrophy in primary myotubes. **a**, Mouse primary myotubes were treated with IKKβ inhibitor TPCA-1 (10 μM), proteosome inhibitor MG132 (5 μM) and EDA-A2 (250 ng/ml) for 24 hr. Changes in gene expression were determined by RT-qPCR (n = 3). **b**,**c**, Primary myotubes were treated with TPCA-1 (10 μM) or proteosome inhibitor MG132 (5 μM) and EDA-A2 (250 ng/ml) for 10 min (**b**) or 24 hr (**c**) and protein levels of NFκB signaling components were studied by western blotting. **d**,**e**, Primary myotubes were treated with recombinant EDA-A2, BAFF or TWEAK proteins (250 ng/ml each) for 24 hr. Protein levels were determined by western blotting (**d**) and gene expression was studied by RT-qPCR (n = 3) (**e**). **f**,**g**, Primary myotubes were transduced with adenoviruses expressing LacZ or human NIK (hNIK). 24hr later, changes in gene expression were determined by RT-qPCR (n = 3) (**f**). Myotubes also treated with GFP adenovirus were visualized 48 hr later under the fluorescence microscope. The scale bar is 50 μm (**g**). The values are mean ± SEM. Statistical analysis was conducted using one-way ANOVA (**a**,**e**) and the two-tailed *t*-test (**f**). \**p* < 0.05, \*\**p* < 0.01, \*\*\**p* < 0.001, compared with the control group or the respective inhibitor only group.

Because EDA-A2 induced mRNA and protein levels of NIK, we compared it with cytokines that are known to activate NIK kinase, such as BAFF and TWEAK^15^. While BAFF failed to trigger an effect in primary myotubes, EDA-A2 and TWEAK acted similarly as both factors stimulated the mRNA expression of the target genes, the accumulation of NIK protein and the processing of p100-NFκB2 (Fig. 2d,e). In fact, TWEAK was previously implicated in muscle atrophy^16,17^. Our findings suggest that both factors may utilize a similar downstream signaling mechanism in myotubes.

If noncanonical NFκB signaling is involved in muscle atrophy, then activation of this pathway should alone stimulate this process. For this purpose, we transduced primary myotubes with an adenovirus expressing NIK. The overexpression of NIK promoted the processing of NFκB2 and the expression of *Atrogin1, MuRF1* and other EDA-A2 targets (Extended Data Fig. 6a and Fig. 2f). In fact, NIK overexpression was sufficient to induce cellular atrophy in primary myotubes (Fig. 2g and Extended Data Fig. 6b). Similar effects on gene expression and cellular atrophy were also observed in human myotubes (Extended Data Fig. 6c-e). We overexpressed mutant mouse NIK isoforms in mouse primary myotubes. While the kinase-dead mutant did not elicit an effect, the autophosphorylation-deficient mutant triggered partial responses in the NFκB2 processing and target gene expression (Extended Data Fig. 6f,g).

We next addressed the necessity of the alternative NFκB activation for EDA-A2-driven myotube atrophy. Treatment of primary myotubes with B022, a specific NIK kinase inhibitor^18,19^, blocked NIK-induced NFκB2 processing and gene expression in a dose-dependent manner (Extended Data Fig. 7a,b). Combined treatment with B022 and recombinant EDA-A2 also inhibited the expression of *Atrogin1, MuRF1* and other EDA-A2 target genes and blunted the NFκB2 processing in primary myotubes (Fig. 3a,b). In fact, the B022 treatment also blocked the EDA-A2-induced mobility shift of NIK protein while the original NIK signal was massively enhanced possibly due to a negative feedback loop broken by the inhibition. A similar effect on NIK protein levels was also observed when EDA-A2-overexpressing primary myotubes were treated with B022 (Extended Data Fig. 7c). Furthermore, overexpression of a dominant-negative NIK form that suppressed the NIK-induced processing of NFκB2 also interfered with the upregulation of atrophy genes by EDA-A2 (Extended Data Fig. 7d-f). Upon NIK kinase inhibition, EDA-A2 was unable to stimulate atrophy in primary myotubes, indicating a major role for NIK signaling in EDA-A2-driven atrophy (Fig. 3c,d).

**Figure 3.**
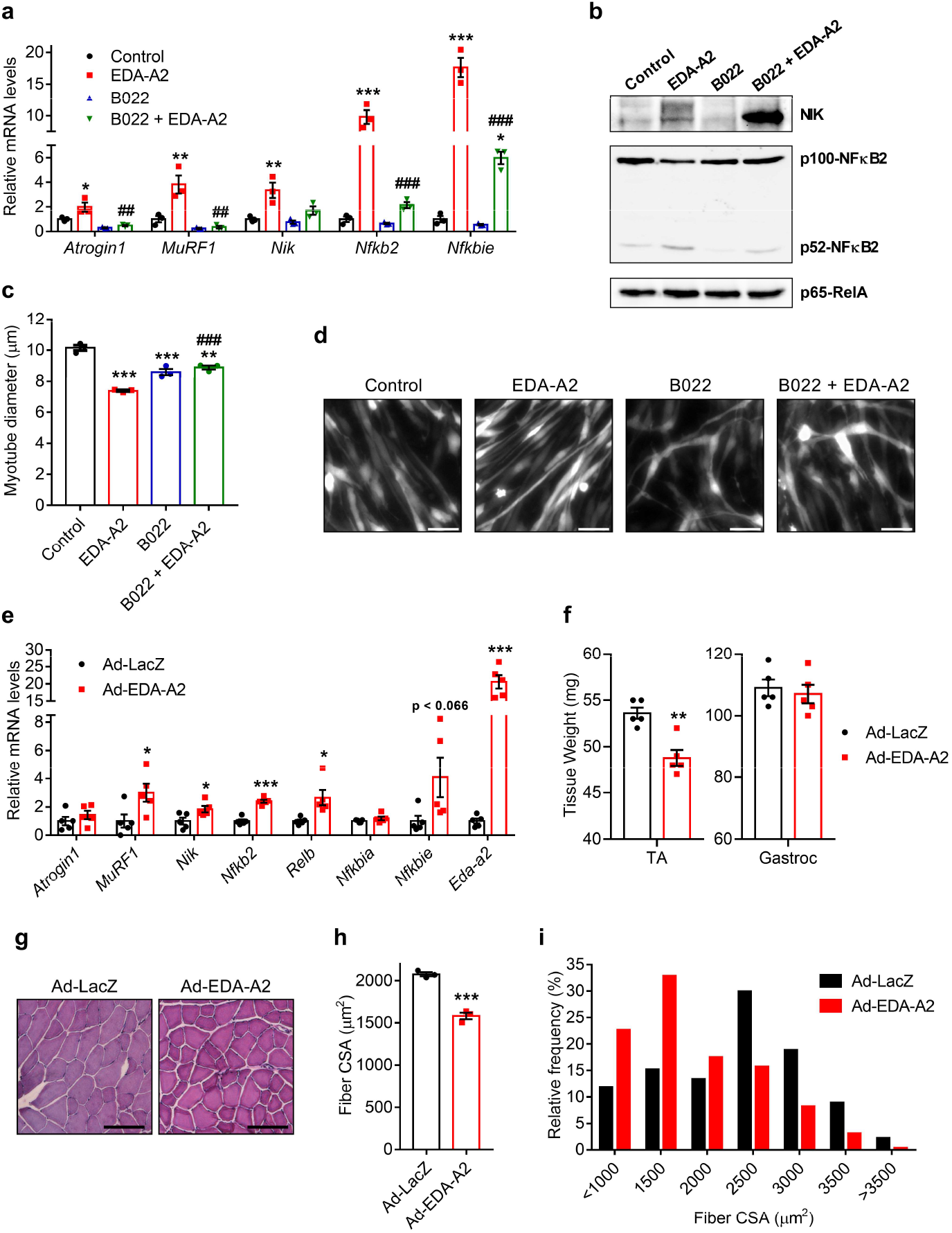
EDA-A2-driven muscle atrophy requires NIK kinase activity. **a**,**b**, Mouse primary myotubes were treated with NIK kinase inhibitor B022 (5 μM) and EDA-A2 (250 ng/ml) for 24 hr. Changes in gene expression were tested by RT-qPCR (n = 3) (**a**) and protein levels were determined by western blotting (**b**). **c**,**d**, Primary myotubes transduced with GFP adenovirus were treated with B022 (5 μM) and EDA-A2 (250 ng/ml) for 48 hr. Cells were visualized under the fluorescence microscope and myotube diameters were measured (n = 3). The scale bar is 50 μm. **e**-**i**, Tibialis anterior muscles of mice were transduced with LacZ or EDA-A2 adenoviruses. Mice were sacrificed 7 days later (n = 5). Changes in gene expression were determined by RT-qPCR (n = 5) (**e**). Tissues were weighed (**f**) and H&E stained (**g**). Muscle fiber cross-sectional area (CSA) (**h**) and the fiber frequency distribution were determined (**i**) (n = 3). The scale bar is 100 μm. The values are mean ± SEM. Statistical analysis was conducted using one-way ANOVA (**a**,**c**) and the two-tailed t-test (**e**,**f**,**h**). \**p* < 0.05, \*\**p* < 0.01, \*\*\**p* < 0.001, compared with the control group or the Ad-LacZ group. *##p* < 0.01, *###p* < 0.001 compares differences between EDA-A2 and B022 + EDA-A2 treatment groups.

EDA-A2’s potent effects on primary myotubes urged us to study its induction in muscle tissue. Previously, transgenic mice overexpressing EDA-A2 in skeletal muscle were generated. These mice exhibited profound muscle degeneration which was prevented by the deletion of EDA2R^8^. Here, we acutely overexpressed EDA-A2 by adenoviral delivery in the tibialis anterior (TA) muscle of mice. Within 7 days, the expression of EDA-A2 target genes, including *MuRF1, Nik, Nfkb2*, and *Relb*, were induced in TA muscles (Fig. 3e). Concomitantly, the weight of TA muscles transduced with EDA-A2 adenovirus significantly dropped while the weight of untreated gastrocnemius muscles remained similar (Fig. 3f). Hematoxylin&eosin (H&E) staining of TA muscles showed that muscle fiber cross-sectional area reduces and the frequency of fibers with small cross-sectional area increases in response to EDA-A2 overexpression (Fig. 3g-i). These results argue that EDA-A2 induction is capable of promoting muscle atrophy *in vivo*.

Next, we addressed the role of EDA2R/NIK signaling in tumor-driven muscle wasting. We utilized EDA2R-null (EDA2R-KO) mice which have normal body weight and lack any obvious phenotypic characteristics^8^. We inoculated littermate wild-type and knockout mice with LLC tumors. Remarkably, muscle wasting was attenuated in EDA2R-KO mice as evidenced by the preservation of gastrocnemius and TA muscles (Fig. 4a). Accordingly, these mice had significantly higher tumor-free body weight compared to tumor-bearing wild-type mice (Extended Data Fig. 8a,b) and they also exhibited improved muscle performance measured by forelimb grip strength (Fig. 4b). Tumor weight, the expression of immune response-related genes in tumors, and plasma C-Reactive Protein (CRP) levels as an indicator of systemic inflammation were comparable between the wild-type and knockout groups (Extended Data Fig. 8b-d). We also dissected and weighed adipose tissue depots, such as epididymal and inguinal white adipose tissue and interscapular brown adipose tissue (BAT). However, a distinct effect on adipose tissue wasting was not detected (Fig. 4a). Similar results on tumor-free body weight, muscle mass and physical strength were obtained when EDA2R-KO mice were inoculated with B16 tumors (Fig. 4c,d and Extended Data Fig. 8e,f).

**Figure 4.**
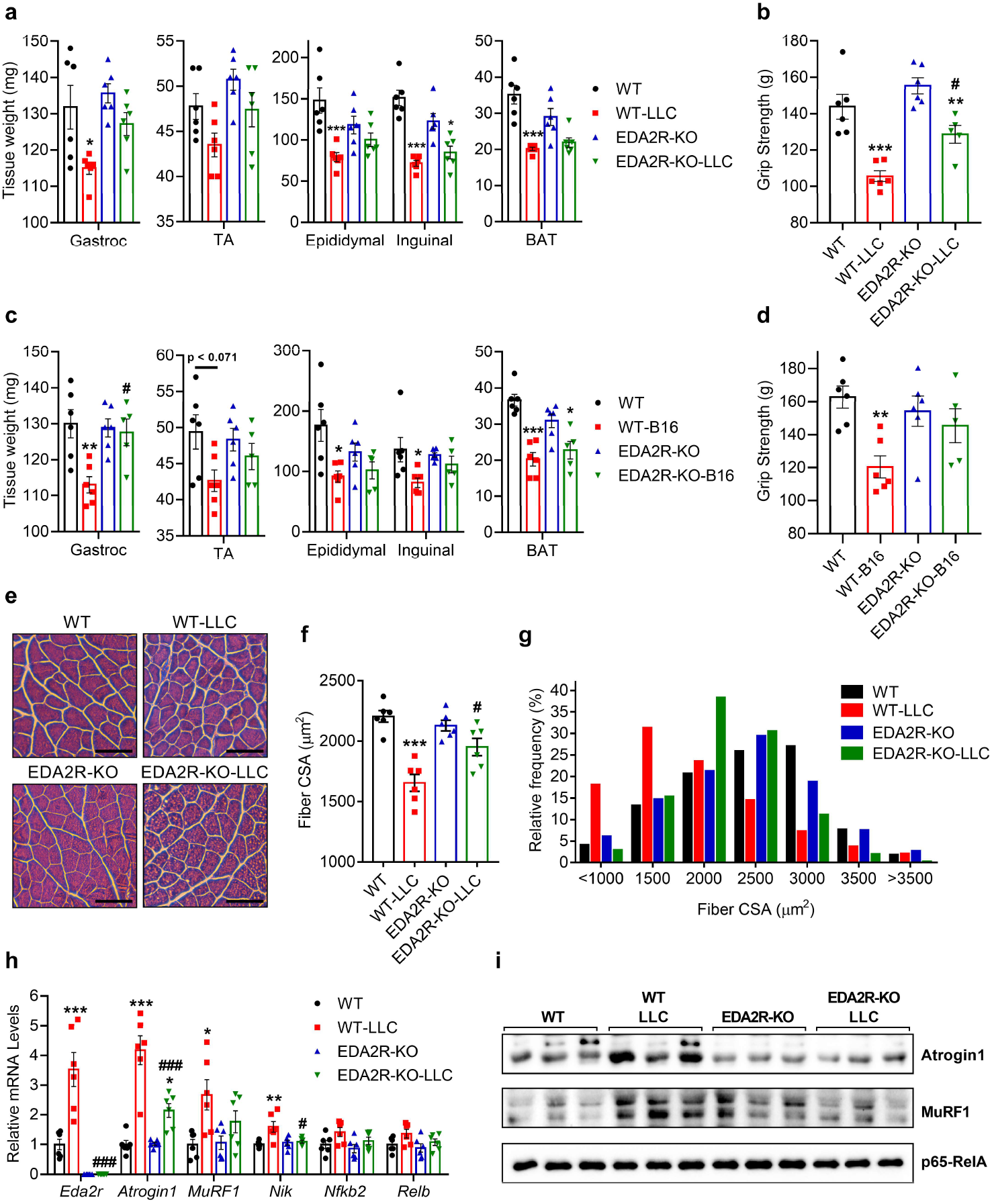
EDA2R-deficient mice are resistant to tumor-driven muscle wasting. **a**,**b**, **e**-**i**, Mice were inoculated with LLC cells and sacrificed 16 days later. Collected tissues were weighed (n = 6) (**a**). Forelimb grip strength was measured before the sacrifice (EDA2R-KO-LLC n = 5, other groups n = 6) (**b**). **c**,**d**, Mice were inoculated with B16 cells and sacrificed 14 days later (EDA2R-KO-B16 n = 5, other groups n = 6). Collected tissues were weighed (**c**). Forelimb grip strength was measured before the sacrifice (**d**). **e**-**g**, Gastrocnemius muscle cross-sections were H&E stained (**e**), cross-sectional area (**f**) and the fiber frequency distribution (**g**) were measured (n = 6). The scale bar is 100 μm. **h**,**i**, Gastrocnemius muscle mRNA levels were tested by RT-qPCR (n = 6) (**h**) and their protein levels were determined by western blotting (n = 3) (**i**). The values are mean ± SEM. Statistical analysis was conducted using two-way ANOVA. \**p* < 0.05, \*\**p* < 0.01, \*\*\**p* < 0.001 compares differences between tumor-bearing and non-tumor-bearing mice of the same genotype. *#p* < 0.05, *###p* < 0.001 compares differences between tumor-bearing wild-type and tumor-bearing knockout mice.

The improvements in muscle mass and function were also reflected in muscle histology. H&E staining of muscle tissue demonstrated an increase in muscle fiber cross-sectional area in tumor-bearing knockout mice compared to wild-type counterparts (Fig. 4e,f, Extended Data Fig. 8g,h). Tumor-driven enrichment of muscle fibers with a small cross-sectional area was suppressed in EDA2R-deficient mice (Fig. 4g and Extended Data Fig. 8i). We also examined changes in the expression of EDA-A2 target genes in these samples. In the absence of EDA2R, tumor-induced mRNA expression of *Atrogin1, MuRF1* and *Nik* was suppressed while a limited induction in *Nfkb2* and *Relb* mRNA levels was detected (Fig. 4h and Extended Data Fig. 8j-l). Furthermore, Atrogin1 and MuRF1 protein levels were also reduced in the muscles of tumor-bearing EDA2R-KO mice (Fig. 4i). These findings indicate that EDA2R function is essential for tumor-driven muscle wasting.

To further delineate this pathway, we generated skeletal muscle-specific NIK knockout mice (Myo-NIK-KO). We confirmed that the deletion was restricted to skeletal muscle by comparing *Nik* mRNA levels in various tissues (Extended Data Fig. 9a). We then inoculated these mice with LLC tumors and studied the cachexia phenotypes. Similar to EDA2R-KO mice, Myo-NIK-KO mice were also resistant to tumor-induced weight loss (Extended Data Fig. 9b-c). The lack of NIK in muscles of tumor-bearing mice prevented muscle loss and also preserved muscle function as determined by forelimb grip strength measurements (Fig. 5a,b). A distinct effect on adipose tissue wasting was not observed (Fig. 5a). The examination of muscle histology of tumor-bearing mice also demonstrated an increase in muscle fiber cross-sectional area and a decrease in the frequency of fibers with small cross-sectional area in Myo-NIK-KO mice compared to wild-type counterparts (Fig. 5c-e). Analysis of gene expression in muscle tissues also demonstrated a reduction in tumor-induced mRNA and protein levels of Atrogin1 and MuRF1 in the knockout mice (Fig. 5f,g and Extended Data Fig. 9d). The similarities in the tumor-driven responses shared by Myo-NIK-KO and EDA2R-KO mice suggest that a common pathway involving EDA2R/NIK acts to promote muscle atrophy.

**Figure 5.**
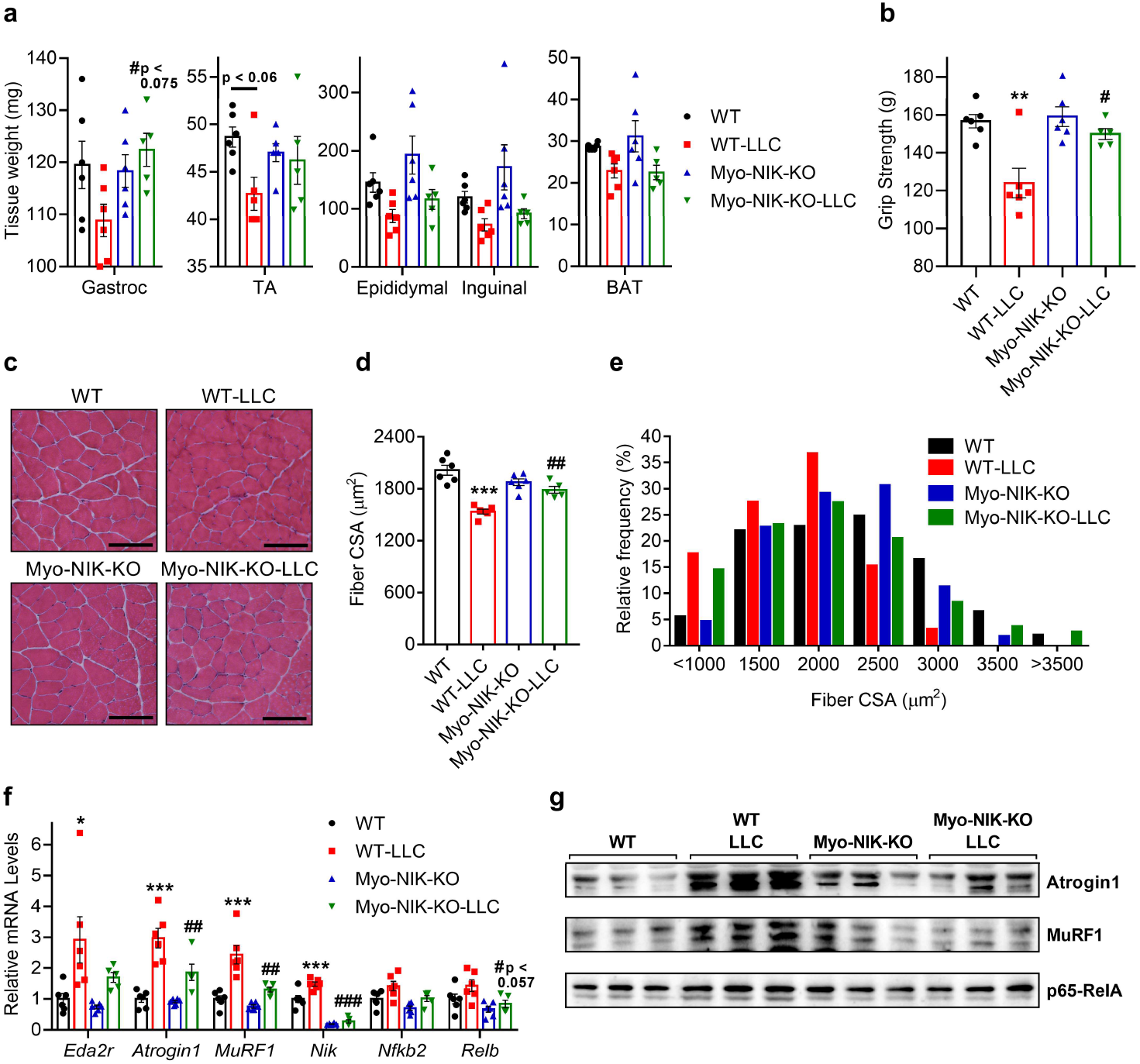
Muscle-specific depletion of NIK protects from tumor-driven muscle wasting. **a**-**g**, Mice were inoculated with LLC cells and sacrificed 16 days later (Myo-NIK-KO-LLC n = 5, other groups n = 6). Collected tissues were weighed (**a**). Forelimb grip strength was measured before the sacrifice (**b**). **c**-**e**, Gastrocnemius muscle cross-sections were H&E stained (**c**), cross-sectional area (**d**) and the fiber frequency distribution (**e**) were measured. The scale bar is 100 μm. **f**,**g**, Gastrocnemius muscle mRNA levels were tested by RT-qPCR (Myo-NIK-KO-LLC n = 5, other groups n = 6) (**f**) and their protein levels were determined by western blotting (n = 3) (**g**). The values are mean ± SEM. Statistical analysis was conducted using two-way ANOVA. \**p* < 0.05, \*\**p* < 0.01, \*\*\**p* < 0.001 compares differences between WT and WT-LLC groups. *#p* < 0.05, *##p* < 0.01, *###p* < 0.001 compares differences between WT-LLC and Myo-NIK-KO-LLC groups.

We also investigated how tumors upregulate *Eda2r* expression in muscle tissue. Testing various tumor-induced cytokines on primary myotubes, we observed *Eda2r* upregulation by Oncostatin M (OSM) (Fig. 6a and Extended Data Fig. 10a), an IL-6 family cytokine involved in a variety of biological processes, including muscle atrophy^20,21^. When overexpressed in TA muscles of mice, OSM significantly increased *Eda2r* mRNA (Fig. 6b). Analysis of blood plasma from LLC tumor-bearing mice detected elevated OSM levels (Fig. 6c), implying that tumor-induced OSM may activate the EDA2R signaling in muscle tissue. In fact, treatment of mouse primary myotubes with recombinant OSM protein also stimulated the expression of *Atrogin1* and resultant cellular atrophy, indicating that OSM itself is an atrophy-inducing factor (Fig. 6d, Extended Data Fig. 10b,c). We found that combined treatment of OSM and EDA-A2 resulted in atrophy-related gene expression in primary myotubes in an additive manner. After testing changes in mRNA levels of OSM target genes; *Osmr* and *Socs3*, EDA-A2-specific targets; *MuRF1, Nik, Nfkb2* and *Relb*, and OSM/EDA-A2 common gene targets; *Atrogin1* and *Ampd3*, we detected additive effects on the expression of *Atrogin1, Ampd3* and *Osmr* (Fig. 6d). Combination of OSM and EDA-A2 also caused a further reduction in myotube diameter (Extended Data Fig. 10b,c). These secreted factors may operate alone or together to induce the atrophy of cultured myotubes.

**Figure 6.**
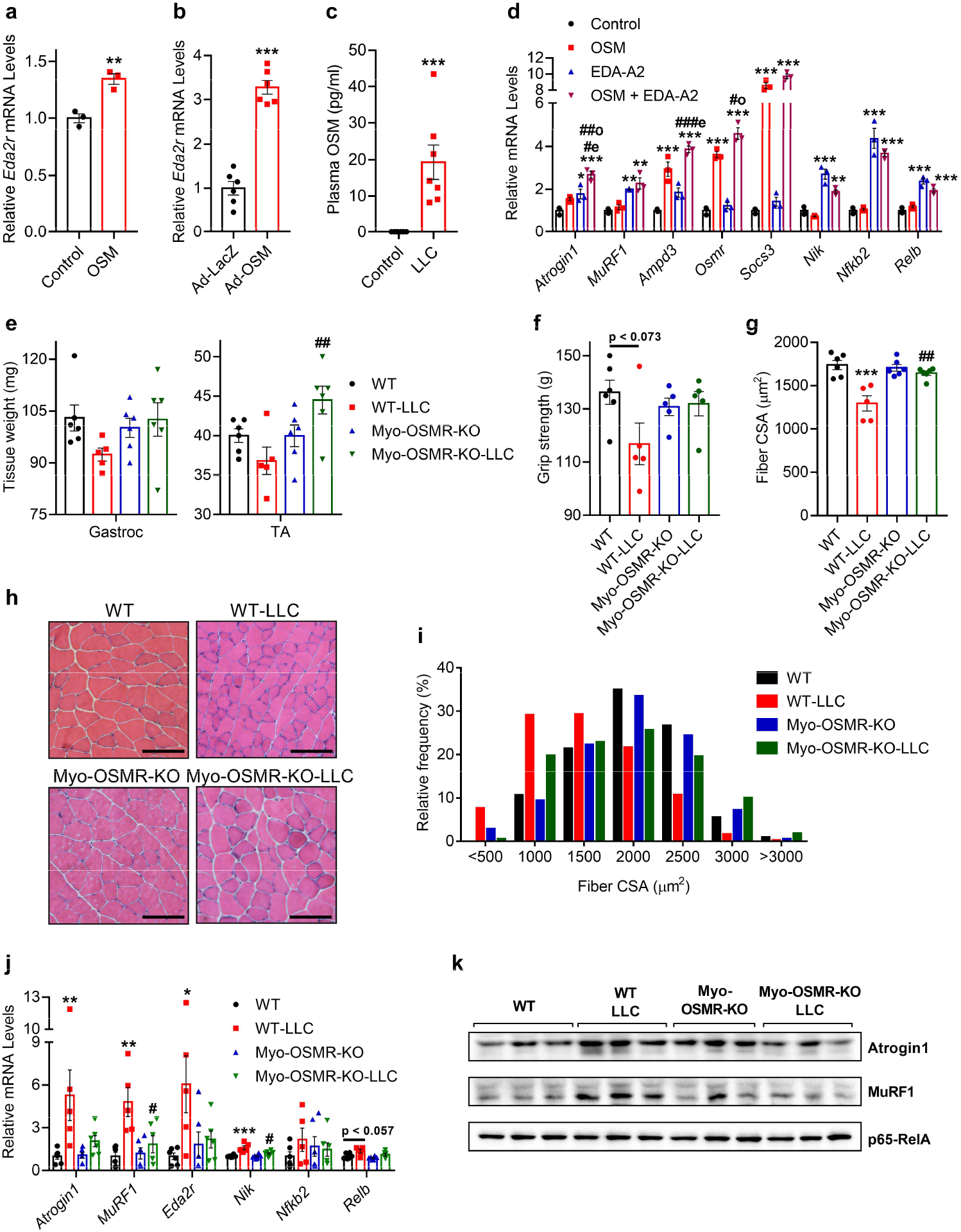
OSM induces *Eda2r* expression in muscle and the depletion of OSMR protects from muscle wasting. **a**, Mouse primary myotubes were treated with recombinant OSM (250 ng/ml) for 6 hr (n = 3). **b**, Tibialis anterior muscles of mice were transduced with LacZ or OSM adenoviruses. Mice were sacrificed 7 days later. mRNA levels were determined by RT-qPCR (n = 6). **c**, Mice were inoculated with LLC cells and sacrificed 16 days later. Plasma OSM levels were measured by ELISA (control n = 8, LLC n = 7) **d**, Mouse primary myotubes were treated with recombinant OSM (250 ng/ml for 48hr) and EDA-A2 (100 ng/ml for 24 hr). Changes in gene expression were determined by RT-qPCR (n = 3). **e-k**, Mice were inoculated with LLC cells and sacrificed 16 days later (WT-LLC n = 5, other groups n = 6). Collected tissues were weighed (**e**). Forelimb grip strength was measured before the sacrifice (WT n = 6, other groups n = 5) (**f**). Gastrocnemius muscle cross-sections were H&E stained (**h**), cross-sectional area (**g**) and the fiber frequency distribution (**i**) were measured (WT-LLC n = 5, other groups n = 6). The scale bar is 100 μm. **j**,**k**, Gastrocnemius muscle mRNA levels were tested by RT-qPCR (WT-LLC n = 5, other groups n = 6) (**j**) and their protein levels were determined by western blotting (n = 3) (**k**). The values are mean ± SEM. Statistical analysis was conducted using the two-tailed t-test (**a**,**b**,**c**), one-way ANOVA (**d**), and two-way ANOVA (**e**,**f**,**g**,**j**). \**p* < 0.05, \*\**p* < 0.01, \*\*\**p* < 0.001, compared to the Control or Ad-LacZ groups (**a**,**b**,**c**,**d**). *#op* < 0.05, *##op* < 0.01, compared to the OSM group, and *#ep* < 0.05, *###ep* < 0.001 compared to the EDA-A2 group (**d**). \**p* < 0.05, \*\**p* < 0.01, \*\*\**p* < 0.001 compares differences between WT and WT-LLC groups and *#p* < 0.05, *##p* < 0.01 compares differences between WT-LLC and Myo-OSMR-KO-LLC groups (**e**,**g**,**j**).

To determine the role of OSM in EDA2R regulation and muscle wasting, we generated skeletal muscle-specific OSM receptor knockout (Myo-OSMR-KO) mice. We confirmed the depletion of OSMR in muscle fibers using immunohistochemistry (Extended Data Fig 10d). Upon LLC tumor inoculation, Myo-OSMR-KO mice were protected from weight loss and muscle wasting (Fig. 6e and Extended Data Fig. 10e,f). However, tumor-bearing knockout mice still lost adipose tissue mass (Extended Data Fig. 10g). Muscle strength assessed by forelimb grip measurements reflected the preservation of muscle mass in the knockout mice while the wild-type mice exhibited evident reduced performance (Extended Data Fig. 6f). H&E staining of gastrocnemius tissue sections indicated that tumor-bearing Myo-OSMR-KO mice had significantly wider muscle fibers than tumor-bearing controls (Extended Data Fig. 6g,h). Fibers with small cross-sectional area were enriched in the latter group (Extended Data Fig. 6i). Gene expression analysis demonstrated that tumor-induced mRNA and protein levels of Atrogin1 and MuRF1 are reduced in muscles of Myo-OSMR-KO mice compared to the wild-type counterparts (Fig. 6j,k and Extended Data Fig. 10h). Importantly, OSMR depletion also suppressed the upregulation of *Eda2r* and its downstream targets in these samples (Fig. 6j and Extended Data Fig. 10h). These findings argue that the OSM/OSMR pathway plays a major role in tumor-driven muscle wasting, including the activation of EDA2R/NIK signaling.

## Discussion

Taken together, our findings indicate that EDA2R signaling is a potent inducer of muscle atrophy. While the EDA-A1/EDAR pathway is important for the development of ectodermal structures, EDA-A2/EDA2R signaling is involved in skeletal muscle pathophysiology^8^. EDA2R expression is upregulated in muscles of tumor-bearing mice and also in muscle biopsies of cachectic cancer patients. Previously, chronic activation of EDA2R signaling was shown to cause myodegeneration in EDA-A2 transgenic mice^8^. Our results show that EDA2R upregulation during cancer cachexia contributes to muscle loss. The canonical NFκB signaling has been shown to promote muscle protein breakdown^2^. This study revealed that noncanonical NFκB activation also triggers muscle atrophy in which NIK kinase plays a central role. Deletion of either EDA2R or NIK in mice was sufficient to confer resistance against tumor-induced muscle wasting. EDA2R expression is induced by inflammatory cytokines and we identified OSM as a prominent regulator. In fact, the depletion of OSMR in muscle protected from the wasting of this tissue. Our results argue that OSM/OSMR signaling acts in parallel to the EDA2R/NIK pathway and reinforces muscle atrophy and EDA2R upregulation.

Systemic inflammation has been implicated to play a major role in cancer cachexia and pro-inflammatory cytokines TNFα, IL-1, and IL-6 were described as causal agents^22^. However, clinical trials testing anti-TNFα therapies failed to prevent muscle atrophy in patients with advanced cancer cachexia ^23,24^. Although anti-IL-1 and anti-IL-6 therapies showed promising results in cachectic cancer patients, a satisfactory effect on skeletal muscle mass was not achieved^25,26^. An effective therapy against cachexia-associated muscle wasting is urgently needed. Our findings argue that EDA2R/NIK and OSM/OSMR pathway elements may serve as novel therapy targets. Prospective studies should test the pharmacologic inhibition of these pathways to prevent muscle wasting. The blockade of EDA2R or OSMR and the inhibition NIK kinase may be useful in reversing muscle loss. Because pathways parallel to EDA-A2/EDA2R, such as TWEAK/FN14, may also utilize NIK to promote muscle atrophy, NIK inhibition stands as a preferable choice. However, the ubiquitous expression of NIK in the body and its prominent roles in immunity may limit this strategy. Interestingly, the depletion of muscle OSMR was sufficient to both attenuate muscle loss and silence the EDA2R activation, making the OSM/OSMR pathway a potentially attractive therapeutic target.

EDA-A2 was previously shown to contribute to obesity-related glucose intolerance possibly through promoting insulin resistance in muscle^14^. Impaired glucose metabolism during cancer cachexia has also been reported^27^. Therefore, it is possible that the inhibition of the EDA2R/NIK pathway may improve cachexia-related abnormalities in glucose metabolism. Furthermore, our gene expression analysis demonstrated elevated *EDA2R* mRNA in the muscle biopsies of muscular dystrophy patients and *EDA2R* upregulation was also reported in muscles of aging individual^28^. It is likely that the role of EDA2R/NIK in muscle atrophy is not restricted to cancer cachexia and therapeutic targeting of this pathway may be beneficial in other muscle disorders.

## Methods

### Reagents

Recombinant proteins were purchased from R&D Systems: EDA-A1 (3944-ED), EDA-A2 (922-ED), TNFα (410-MT), BAFF (8876-BF), TWEAK (1237-TW), OSM (495-MO), IL-6 (406-ML) and LIF (8878-LF). Small molecule inhibitors were acquired from indicated sources: TPCA-1 (abcam, ab145522), MG132 (Sigma; M8699), BAY 11-7082 (abcam, ab141228), BOT-64 (Santa Cruz; sc-222062), B022 (Aobious; AOB8699). Mouse EDA-A1 (MC208411), mouse EDA-A2 (MC208415) and mouse OSM (MR226014) expression plasmids were purchased from Origene. Mouse NIK expression plasmid was purchased from Invivogen (pUNO1-mMap3k14). Wild-type human NIK and the NIK-K429/430A mutant plasmids were a gift from Prof. Michael Kracht (JLU Giessen).

### Mice

Mice were housed in 12 hour light/dark cycles (7am-7pm) and given ad libitum access to a standard rodent chow diet and water. 8-12-week-old male mice were used in all animal experiments. Mice were kept in the Koc University Animal Research Facility in accordance with institutional policies and animal care ethics guidelines. EDA2R-KO and NIK-floxed mice were generated by Genentech^8,29^ NIK-floxed mice were a gift from Prof. Shao-Cong Sun (MD Anderson Cancer Center). Myo-NIK-KO mice were generated by crossing NIK-floxed and ACTA1-Cre mice (Jackson strain #006149). OSMR-floxed mice (strain #011081) were purchased from Jackson laboratory. Myo-OSMR-KO mice were generated by crossing OSMR flox and ACTA1-Cre mice (Jackson strain #006149). All mice were maintained on a pure C57BL/6 background. Plasma CRP levels were measured using an ELISA assay (BT Lab E0218Mo). All animal protocols were approved by the Institutional Animal Care and Use Committee of Koc University.

### Tumor inoculation

LLC and B16 (B16-F10) cells were cultured in DMEM medium (Sigma 5796) with 10% Fetal Bovine Serum (FBS) and penicillin/streptomycin (Invitrogen). B16 cells were also supplemented with freshly added 2 mM L-glutamine (Invitrogen). Mice were divided into groups randomly while satisfying the criteria that the average body weight in each group is similar. All mice used in tumor inoculation experiments including the transgenic lines were from C57BL/6 background. LLC cells (5 × 10^6^ per mouse) or B16 (2.5 × 10^6^ per mouse) cells were injected subcutaneously over the flank. Non-tumor-bearing control mice received the vehicle (PBS) only. Mice were housed individually in all tumor inoculation experiments. Mice were sacrificed 16 days (LLC) or 14 days (B16) post tumor inoculation. Epididymal, inguinal and interscapular fat depots, gastrocnemius and tibialis anterior muscles and tumors were dissected and weighed using an analytical balance.

### Grip strength

Forelimb grip strength was measured on the same day as the sacrifice. Each mouse was allowed to grab a bar attached to a force transducer while the mouse was steadily pulled by the tail horizontally away from the bar (Ugo Basile grip strength meter). The maximum strength produced before releasing the bar registered from at least 3 repetitions was averaged to determine the grip strength of each mouse.

### Tissue histology

Isopentane was cooled by liquid nitrogen until 2/3 is frozen. Muscle samples wrapped in aluminum foil were placed in the cooled isopentane for 15-20 seconds. Frozen tissues were embedded in Tissue-Tek OCT freezing medium (Sakura) in cryomolds. Using a cryostat, 8 μm thick sections were cut and collected on Superfrost Plus slides (Thermo). Sections were fixed with 4% paraformaldehyde and treated with hematoxylin (Merck 105174), 0.1% HCl, eosin (Merck 109844), 70-100% ethanol gradient and xylene (Isolab), respectively. Muscle fiber cross-sectional area was measured using Image J software.

### Immunohistochemistry

5 μm thick cryosections were fixed with neutral buffer formalin and then incubated with 0.3% hydrogen peroxide for 10 min to block endogenous peroxidase activity. Sections were incubated in blocking solution (3% bovine serum albumin + 0.1% Triton X-100 + 5% Horse serum) at room temperature for 1hr and then with anti-OSMR beta (R&D Systems, AF662) antibody (10 μg/ml) in blocking solution overnight at +4°C. Sections were washed with PBS and incubated in the blocking solution containing anti-goat IgG H&L HRP-conjugated secondary antibody (1:1000) (Abcam, ab6885) for 1hr. Finally, the sections were stained with diaminobenzidine (DAB; Abcam, ab64238,) and counterstained with hematoxylin (Merck 105174).

### Adenovirus production and injection

Adenovirus vectors were generated using the Virapower Adenoviral expression system (Invitrogen). Briefly, open reading frames of the genes following a CACC sequence were cloned into a pENTR-D-TOPO plasmid and then recombined into a pAd-CMV-DEST adenoviral plasmid using LR clonase II. PacI (Thermo) digested adenoviral plasmids were transfected into 293A cells using Lipofectamine 2000 (Invitrogen). 293A cells were cultured in DMEM (Sigma 5796), 10% FBS (Invitrogen) and penicillin/streptomycin (Invitrogen). Adenoviral particles were collected from cell culture supernatant following the manufacturer’s instructions. For purification and titration, Adeno-X Maxi Purification Kit and Adeno-X Rapid Titer Kit from Clontech were used. Mice were injected unilaterally with 5×10^8^ ifu (infectious units) of Adeno-EDA-A2 or Adeno-OSM into the tibialis anterior muscle while the contralateral muscle received the same dose of control Adeno-LacZ. Mice were sacrificed 7 days later.

### Site-directed mutagenesis

Mouse NIK coding sequence in the pENTR shuttle plasmid was mutated using Phusion high fidelity polymerase (NEB) and oligonucleotides designed to carry the desired mutations (indicated by lowercase letters): mNIK-T561A F: 5’-CTACATTCCTGGCgCGGAGACCCACATG-3’, R: 5’-CATGTGGGTCTCCGcGCCAGGAATGTAG-3’. mNIK-K431A-K432A 5’-GCTTCCAGTGTGCTGTCgcAgcGGTACGACTCGAGGTG-3’ R: 5’-CACCTCGAGTCGTACCgcTgcGACAGCACACTGGAAGC-3’. Mutated coding sequences were recombined into the adenoviral plasmid for adenovirus production as described above.

### Myoblast culture

Primary myoblasts were isolated from limb muscles of mice (2-3 days old) as described before^30^. Myoblasts were cultured in Ham’s F-10 nutrient mixture (Invitrogen) with 20% FBS (Invitrogen) supplemented with 2.5 ng/ml basic fibroblast growth factor (bFGF) (Sigma) and penicillin/streptomycin (Invitrogen). For differentiation, myoblasts were then transferred to DMEM (Sigma 5796) supplemented with horse serum (HS) and penicillin/streptomycin (Invitrogen). Myotube cells were differentiated in 2% HS for 48 hours for protein isolation and they were differentiated in 5% HS for 72 hours for gene expression and imaging experiments. Adeno-GFP was added to the cells at the start of differentiation for fluorescent myotube imaging performed using a live cell imager (Zeiss Axiolab live cell imager). Cells were treated with other adenoviruses after differentiation for 24-48 hr. Human Skeletal Muscle Myoblasts (HSMM; Lonza) were cultured in DMEM/F12-glutamax (Invitrogen) with 20% FBS (Invitrogen) supplemented with 5 ng/ml bFGF (Sigma). Cells were differentiated in DMEM/F12-glutamax with 5% HS for 72 hours. Human myotubes were treated with recombinant proteins or adenoviruses for 48 hr. The diameters of individual myotubes were measured using Image J software. Each myotube was measured at 3 different sites and the values were averaged.

### Immunofluorescence

Cells were fixed with ice-cold 100% methanol at −20°C for 10 min, incubated in a blocking solution (3% bovine serum albumin + 0.1% Triton X-100 + 10% Horse serum) at room temperature for 1hr and then incubated with myosin heavy chain (MyHC) antibody (1:1000) (DSHB, MF20) in the blocking solution for 1 hr at room temperature. Cells were washed with PBS and incubated with anti-mouse IgG H&L Alexa Fluor 594 secondary antibody (1:2000) (Abcam, ab150116) and DAPI (1:3000) (Cayman, 14285) in the blocking solution for 1hr and mounted using homemade mounting medium. Cells were visualized using fluorescence microscopy (Zeiss). MyHC signal was normalized to the number of myotube nuclei. Fields with similar density of myotubes were chosen.

### Western blotting

Cells were homogenized in a cell lysis buffer containing 50 mM Tris (pH 7.4), 150 mM NaCl, 1% Triton X-100, 5 mM EDTA, 1 mM PMSF, supplemented with protease inhibitor tablets (Roche) and phosphatase inhibitors; 20 mM NaF, 10 mM β-glycerol phosphate, 10 mM Na_4_P_2_O_7_, 2 mM Na_3_VO_4_. A similar lysis buffer was used for tissue samples where 1% NP40 was used as the detergent and 10% glycerol was added. Tissues were homogenized using a Kinematica (PT1200E) homogenizer. The homogenates were centrifuged at 13,000 rpm for 10 min and the supernatants were used as lysates. Protein concentration was determined by Bio-Rad Protein assay and 30 μg of protein lysate was used in each SDS-PAGE run. For the detection of endogenous NIK protein and its mobility shift, 90 μg of protein lysate was loaded. Nitrocellulose membrane was blotted with primary antibodies in TBS containing 0.05% Tween and 5% BSA (Cell signaling; p65-RelA (8242), phospho-p65-RelA-Ser536 (3033), p105/p50-NFκB (12540), phospho-p105-NFκB-Ser932 (4806), p100/p52-NFκB2 (4882), IκB (4814), phospho-IκB-Ser32 (2859), NIK (4994), JNK (9252), phospho-JNK-Thr183/Tyr185 (4668), anti-Flag/DYKDDDDK (2368), ECM Bioscience; Atrogin1 (AP2041), MuRF1 (MP3401), and DSHB; MyHC antibody (MF20)). For secondary antibody incubation, TBS-T containing 5% milk was used (Cell signaling anti-rabbit (7074), anti-mouse (7076)). WesternBright blotting substrates from Advansta were used for visualization of the results on a Chemidoc imaging system (Bio-rad). For blots visualized with a Licor Odyssey CLx imaging system, IRDye 680RD anti-mouse (926-68070) and IRDye 800CW anti-rabbit (926-32211) secondary antibodies were used.

### In vitro phosphatase assay

A cell lysis buffer similar to the recipe described above but lacking EDTA and phosphatase inhibitors was used to collect cells. The protein concentration of the homogenate supernatants was determined by Bio-Rad Protein assay. 90 μg of lysate was mixed with or without alkaline phosphatase (0.1U/μg lysate) and its buffer (Thermo) and incubated at 37°C for 1 hour. Samples were run on SDS-PAGE and processed as described above.

### RT-qPCR

Total RNA from cultured cells or tissue samples was extracted using Qiazol reagent (Qiagen) and purified with RNA spin columns (Ecotech). Tissues were homogenized using TissueLyzer LT (Qiagen). Complementary DNA synthesis was carried out with a High-Capacity cDNA Reverse Transcription kit (Thermo). The resultant cDNA was analyzed by RT–qPCR using a CFX Connect instrument (Bio-Rad). In each reaction, 25 ng of cDNA and 150 nmol of each primer were mixed with iTaq Universal SYBR Green Supermix (Bio-Rad). Relative mRNA levels were calculated by the ΔΔCt method and normalized to cyclophilin mRNA. The following primers were used: *Cyclo*F: 5’-GGAGATGGCACAGGAGGAA-3’, R: 5′-GCCCGTAGTGCTTCAGCTT-3′. *Atrogin1* F: 5′-TCAGAGAGGCAGATTCGCAA-3′, R: 5′-GGGTGACCCCATACTGCTCT-3′. *MuRF1* F: 5′-TCCTGATGGAAACGCTATGGAG-3′, R: 5′-ATTCGCAGCCTGGAAGATGT-3′. *Eda2r* F: 5′-TCCCCTCTACTGGACCTGAA-3′, R: 5′-TGAAAGAGACCTTTCTAGTTCACCT-3′. *Eda2r-*KO F: 5′-CAGGACCAAGAATGCATCCCA-3′, R: 5′-GCTCAACTGGAAGGTACACTGAA-3′. *Eda-a2* F: 5′-TCAAAAATGATCTTTCAGGTGGAG-3′, R: 5′-TGAAGTTGATGTAGTAGACCTG-3′. *Edar* F: 5′-TTGTTGAAGGTCTCAGCCCC-3′, R: 5′-TTTTCACGACCGCCTTCTCA-3′. *Eda-a1* F: 5′-TCTTTCAGGTGGAGTGCTCA-3′, R: 5′-TGAAGTTGATGTAGTAGACTTCTAC-3′. *Nik* F: 5′-CGAGCTACTTCAACGGGGTC-3′, R: 5′-GGCAATGTCTCCCACCTTGA-3′. Nik-KO F: 5′-TGTTCTGTGGGAAGTGGGAG −3′, R: 5′-CTCTTGGCTATTCTCACATTCAGC-3′. *Nfkb1* F: 5′-CTGAACAATGCCTTCCGGCT-3′, R: 5′-TGGTACCCCCAGAGACCTCAT-3′. *Nfkb2* F: 5′-CCTTCGTAGTTACAAGCTGGC-3′, R: 5′-GGCACTGTCTTCTTTCACCT-3′. *Nfkbia* F: 5′-TAGCAGTCTTGACGCAGACC-3′, R: 5′-CGTGTGGCCATTGTAGTTGG-3′. *Nfkbib* F: 5′-ACCTCAATAAACCGGAGCCTAC-3′, R: 5′-CACCGGCTTTCAGGAGAAGTT-3′. *Nfkbid* F: 5′-CAGTCATACAAGCCAGGAGAT-3′, R: 5′-TCATATTAACAAAGGCCCGCA-3′. *Nfkbie* F: 5′-GACATTGATGTACAGGAGGGCA-3′, R: 5′-GGTGTGCACCCGTTAAGCAT-3′. *Nfkbiz* F: 5′-CAGTGGAGGCAAAGGATCGTA-3′, R: 5′-GGCAACTCCAAAAAGAGGCG-3′. *Rela* F: 5′-GATCGCCACCGGATTGAAGA-3′, R: 5′-GGGGTTCAGTTGGTCCATTG-3′. *Relb* F: 5′-TTCAAAACGCCACCCTACGA-3′, R: 5′-ACACCGTAGCTGTCATGATCC-3′. *Relc* F: 5′-ATTTATGACAACCGTGCCCCA-3′, R: 5′-CCCTGACACTTCCACAGTTCT-3′. *Ampd3* F: 5′-CTCCTCTCAGCAACAACAGCC-3′, R: 5′-CTCCATGAGCGCTTCCTTTGTG-3′. *Osmr* F: 5′-GGTCCTTCATCCAGCCTTCC-3′, R: 5′-GCTCCTCCAAGACTTCGCTT-3′. *Socs3* F: 5′-TAGACTTCACGGCTGCCAAC-3′, R: 5′-CGGGGAGCTAGTCCCGAA-3′. *Tnfa* F: 5′-CCACCACGCTCTTCTGTCTA-3′, R: 5′-CCATTTGGGAACTTCTCATCCC-3′. *Il6* F: 5′-CACTTCACAAGTCGGAGGCT-3′, R: 5′-TGCCATTGCACAACTCTTTTCT-3′. *Ifng* F: 5′-CTTCAGCAACAGCAAGGCG-3′, R: 5′-CTGTGGGTTGTTGACCTCAAACT-3′. *Il1b* F: 5′-AAGGAGAACCAAGCAACGACA-3′, R: 5′-TTGGGATCCACACTCTCCAGC-3′. *Il10* F: 5′-GTAGAAGTGATGCCCCAGGC-3′, R: 5′-GGGGAGAAATCGATGACAGC-3′. *Ccl2* F: 5′-CACTCACCTGCTGCTACTCA-3′, R: 5′-GCTTGGTGACAAAAACTACAGC-3′. *Ccl5* F: 5′-TGCCCACGTCAAGGAGTATT-3′, R: 5′-TTCGAGTGACAAACACGACTG-3′. *F4/80* F: 5′-CTTCTGGGGAGCTTACGATGG-3′, R: 5′-GGCCAAGGCAAGACATACCA-3′. *Cd68* F: 5′-ACTTCGGGCCATGTTTCTCT-3′, R: 5′-GGGGCTGGTAGGTTGATTGT-3′. *Nos2* F: 5′-CAGGAGATGGTCCGCAAGAG-3′, R: 5′-GTCCTGAACGTAGACCTTGGG-3’. *Arg1* F: 5′-CGTAGACCCTGGGGAACACTAT-3′, R: 5′-TCCATCACCTTGCCAATCCC-3′. *Cd163* F: 5′-GTGTTCCGAAGGACAGGTGG-3′, R: 5′-AAGCTGGCCACTTGCTATGC-3′. *Cd19* F: 5′-GAAGCATCCTCGCTTGGGTC-3′, R: 5′-ACTGGGACCGGACTGAATTG-3′. *Cd3e* F: 5′-TGTATCACTCTGGGCTTGCTG-3′, R: 5′-CTCCTTGTTTTGCCCTCTGGG-3′. *hCYCLO* F: 5′-GGAGATGGCACAGGAGGAA-3′, R: 5′-GCCCGTAGTGCTTCAGTTT-3′. *hNIK* F: 5′-GGGACGTCAAAGCTGACAAC-3′, R: 5′-GACACACAGCATGGCCAAAG-3′. *hEDA2R* F: 5′-TGCCTCCTATACTGGAGCTGA-3′, R: 5′-GGGGCCCAAGAGACCTCATTA-3′. hATROGIN1 F: 5′-AGGAAGTACTAAAGAGCGCCA-3′, R: 5′-GCAGGCCGGACCACGTA-3′. hMURF1 F: 5′-GCCCCATTGCAGAGTGTCTT-3′, R: 5′-ACTGTTCTCCTTGGTCACTCG-3′. hNFKB2 F: 5′-TCTGCAACTGAAACGCAAGC-3′, R: 5′-CCTCTTCCTTGTCTTCCACCA-3′. hRELB F: 5′-CTCGCGACCATGACAGCTAC-3′, R: 5′-GGCTTTTTCTTCCGCCGTTT-3′.

### Human gene expression analysis

Patients were enrolled in the ACTICA study, a cross-sectional study aimed at assessing cachexia in patients diagnosed with colorectal or lung cancer. The study was performed at the Cliniques Universitaires Saint-Luc, Brussels, Belgium from January 2012 to March 2014. The study protocol was approved by the local ethical committee of the Université Catholique de Louvain (protocole code: 2011/19AVR/157, approved on the 9 May 2011) and written consent was given prior to entry into the study. The inclusion and exclusion criteria have been precisely described previously in the original paper^31^. Cachexia was defined, according to the definition proposed by Fearon et al^32^, as an involuntary weight loss >5 % over the past 6 months or weight loss >2% and BMI <20 kg/m^2^ or weight loss >2% and low muscularity. Among 152 patients enrolled in the study, a skeletal muscle microbiopsy was performed in 35 patients under general anesthesia, just before the surgery for cancer and before any other therapeutic intervention. The microbiopsies were taken from the vastus lateralis of the quadriceps, with a 14 Gauge true-cut biopsy needle (Bard Magnum Biopsy gun; Bard, Inc.). The muscle samples were cleaned of gross blood contamination and fat or fibrous tissue prior to being frozen in liquid nitrogen and stored at −80°C until RNA extraction. Total RNA was extracted from frozen muscle samples using TriPure Isolation Reagent (Roche Diagnostics, Basel, Switzerland), as described by the manufacturer. Reverse transcription was performed as previously described by Gueugneau et al^33^.

Gene expression profiles of rectus abdominis muscle biopsies from cachectic and non-cachectic pancreatic ductal adenocarcinoma patients and non-cancer subjects were accessed from Gene Expression Omnibus (GEO) database with the GSE130563 accession number^34^. Differential gene expression analysis was performed by GEO2R with default settings. One sample from GSE130563 was identified as an outlier (GSM3743567) for EDA2R expression according to Grubbs’ test (Alpha = 0.0001) and excluded from the analysis. Gene expression profiles of quadriceps muscle biopsies collected from non-cancer subjects and cachectic patients with upper gastrointestinal cancer before and after the tumor resection were accessed from the GEO database (accession number: GSE34111)^35^ and analyzed by GEO2R using default settings. Gene expression profiles of skeletal muscles samples from DMD patients and normal subjects were accessed with the GSE1007 accession number^36^. Samples grouped as DMD and Normal were analyzed by GEO2R using default settings. RNA sequencing results of muscle biopsy samples from FSHD subjects were also accessed from the GEO database (accession number: GSE115650)^37^. 1-year follow-up gene expression assessment of muscle biopsies from the same patients was also analyzed (accession number: GSE140261)^38^. Gene counts normalization and fold change calculations were performed using the DEseq2 (v1.34.0) R package.

### Statistical Analysis

Values are expressed as mean ± SEM. Error bars (SEM) shown in all results were derived from biological replicates. Significant differences between two groups were evaluated using a two-tailed, unpaired t-test. Comparisons of more than two groups were performed using one-way or two-way ANOVA and corrected for multiple comparisons using Tukey’s post-hoc test. Values of *p* < 0.05 were considered statistically significant. Exact *p*-values and the type of statistical test used in each experiment can be found in the Figure legends.

## Acknowledgements

The authors gratefully acknowledge the use of the animal facility infrastructure at Koc University Research Center for Translational Medicine (KUTTAM). The authors thank Prof. Michael Kracht (JLU Giessen) for sharing human NIK plasmids and Prof. Shao-Cong Sun (MD Anderson Cancer Center) for sharing NIK-floxed mice. EDA2R-null and NIK-floxed mice were provided by Genentech. The authors also appreciate the assistance received from Ibrahim Oguz. D.H.A. was funded by a TUBITAK-BIDEB scholarship. This work was supported by the Scientific and Technological Research Council of Turkey (TUBITAK) grants 118Z167, 118Z791 and 118C014, and the EMBO Installation Grant 4162 to S.K.

## Author contributions

S.K. conceived and designed the experiments. S.N.B., A.D., B.T., S.A., B.Z.C.W., D.H.A., Z.O., P.L., J.P.T., A.L. and S.K. performed the experiments. S.N.B., A.D., B.T., S.A. and S.K. analyzed the data. S.N.B., A.D. and S.K. wrote the manuscript.

## Data Availability

Human gene expression datasets analyzed in this study are available in the GEO database; GSE130563^34^, GSE34111^35^, GSE1007^36^, GSE115650^37^, and GSE140261^38^. A detailed description of the cancer patient muscle biopsies used in this study was previously published^31^.

## Materials & Correspondence

Correspondence and requests for materials should be addressed to S.K.

---

**Extended Data Figure 1.**
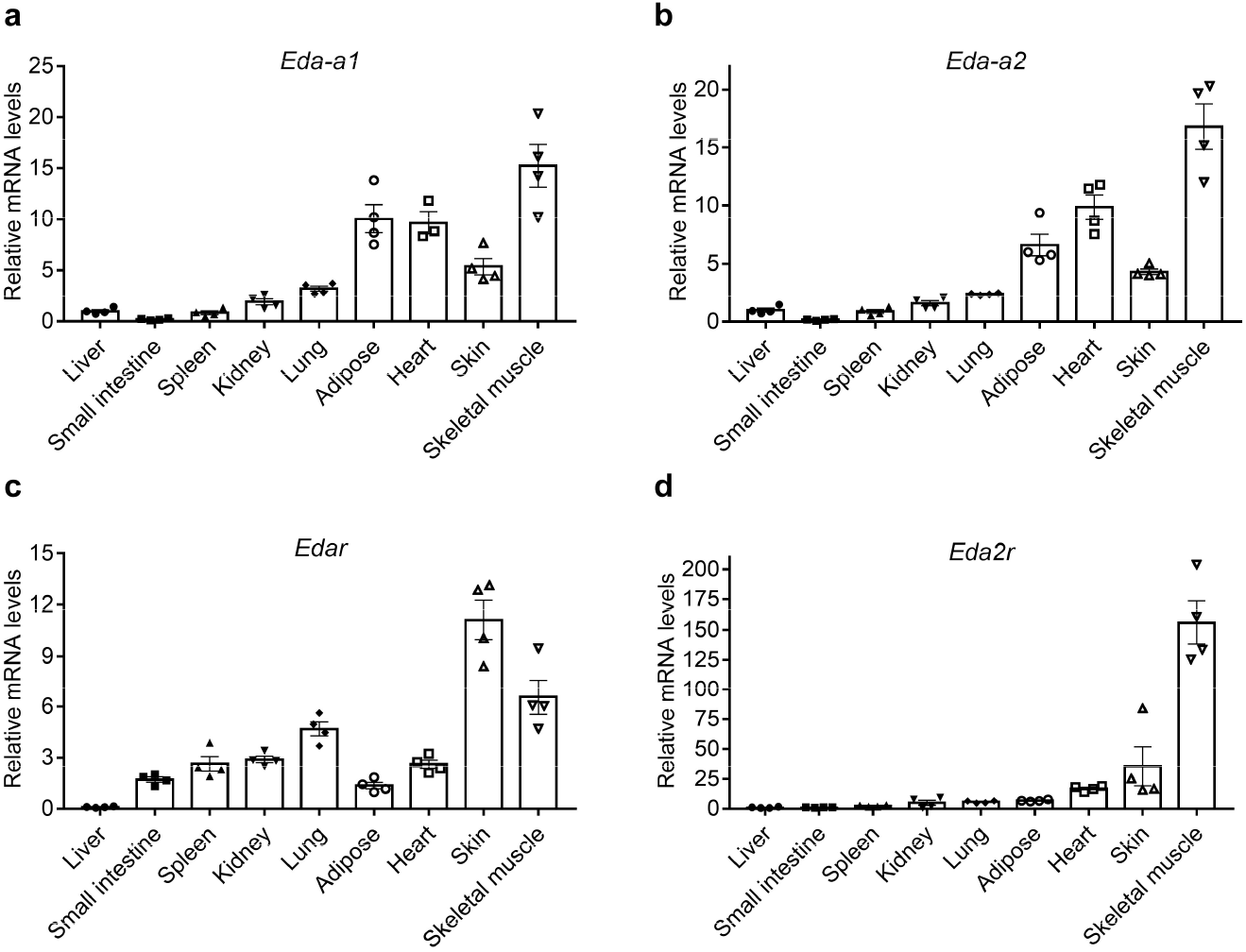
The expression of *Eda-a2* and *Eda2r* is enriched in skeletal muscle tissue. **a**-**d,** Various tissue samples were collected from C57BL/6 mice. Relative mRNA levels were determined by RT-qPCR (n = 4). The values are mean ± SEM.

**Extended Data Figure 2.**
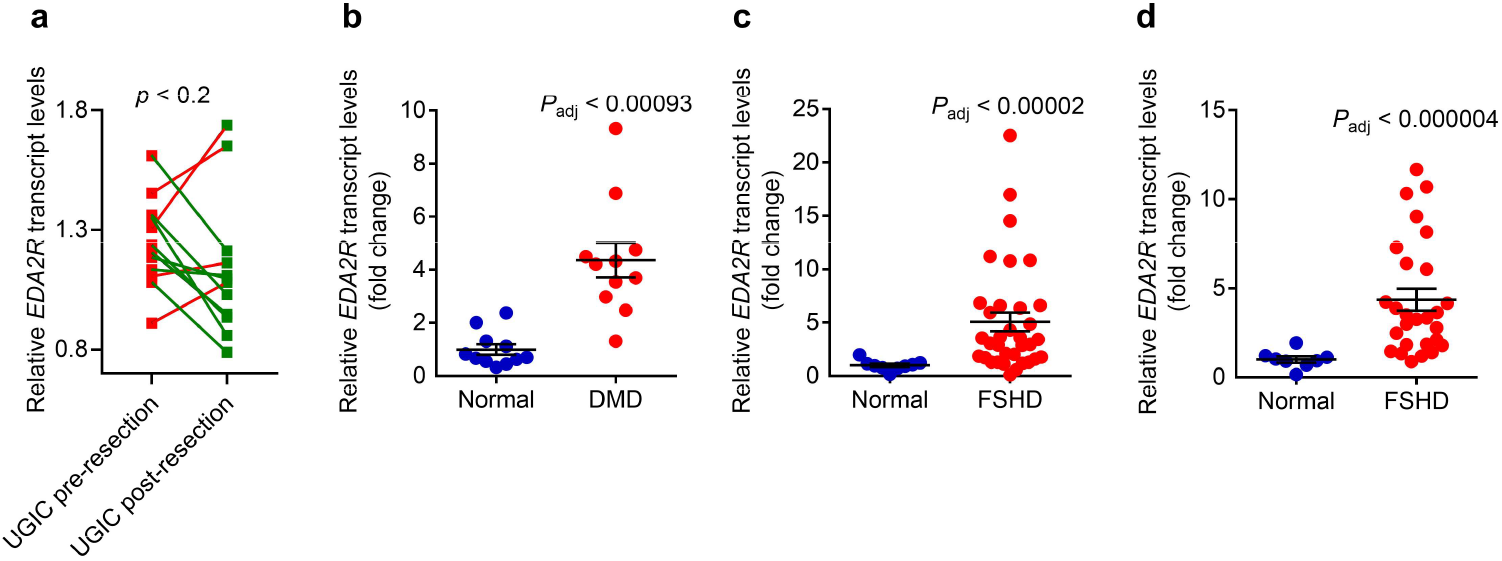
*EDA2R* expression is induced in DMD and FSHD patients. **a**, *EDA2R* transcript levels were analyzed in quadriceps muscle biopsies collected from cachectic patients with upper gastrointestinal cancer (n = 12) before and after tumor resection (GSE34111). Upon surgery, EDA2R levels were upregulated in 4 patients (red connecting lines) and downregulated in 8 patients (green connecting lines). Statistics by two-tailed paired t-test. **b**, GSE1007 dataset was analyzed by GEO2R and *EDA2R* expression values were determined in normal subjects and DMD patients (n = 10). In each group, one individual had 2 technical replicates making the total number of data points 11. **c**, GSE115650 dataset was analyzed by DESeq2 and *EDA2R* expression values were determined in normal subjects (n = 9) and FSHD patients (n = 34). **d**, GSE140261 dataset was analyzed by DESeq2 and *EDA2R* expression values were determined in normal subjects (n = 8) and FSHD patients (n = 27). The values are mean ± SEM. Statistical values were adjusted for multiple tests.

**Extended Data Figure 3.**
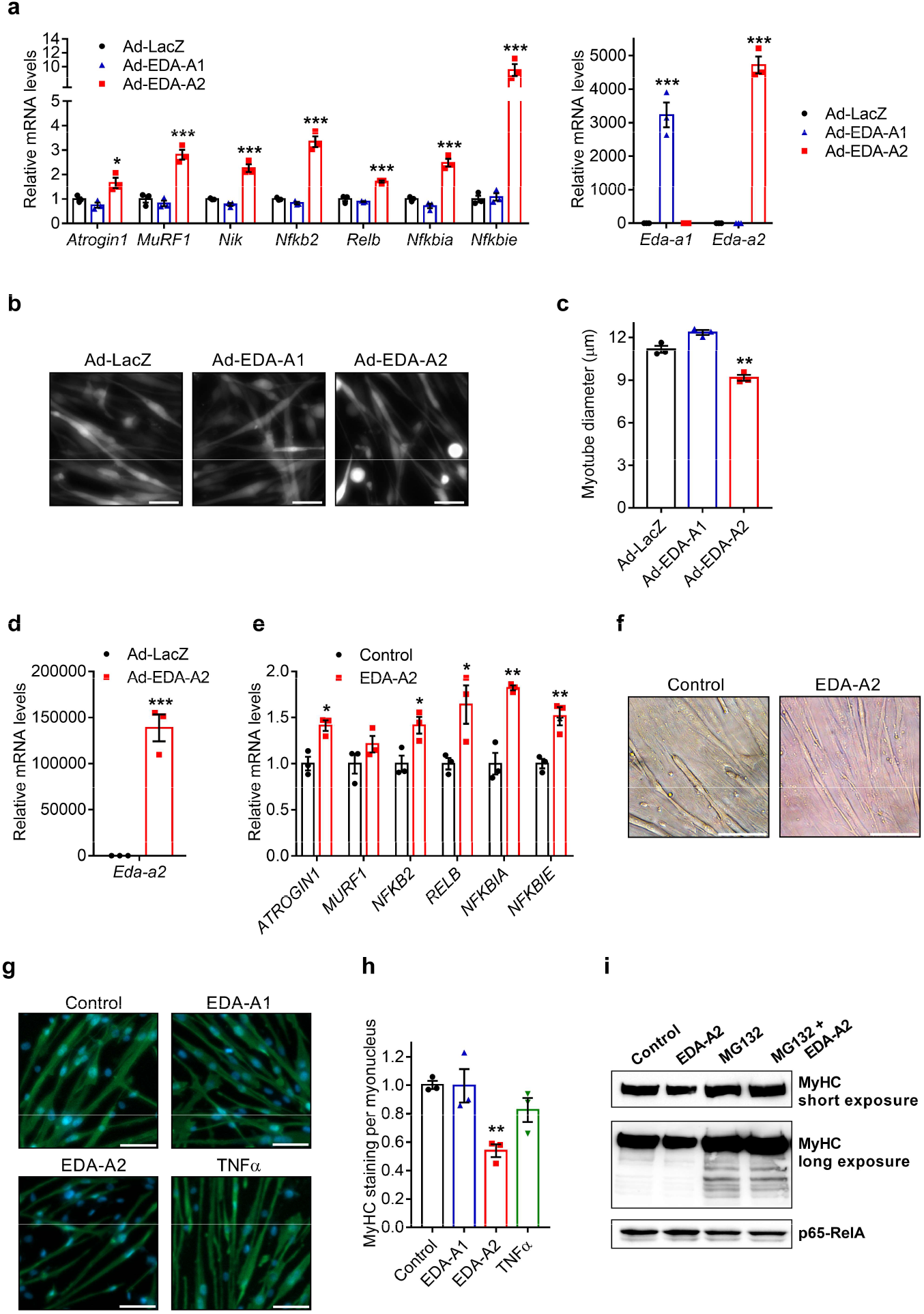
The overexpression of EDA-A2 or the administration of recombinant EDA-A2 in human and mouse myotubes stimulates cellular atrophy. **a**-**c**, Mouse primary myotubes were transduced with LacZ, EDA-A1, or EDA-A2 expressing adenoviruses. 24 hr later, gene expression was tested by RT-qPCR (n = 3) (**a**). Myotubes also treated with GFP adenovirus were visualized 48 hr later under the fluorescence microscope. The scale bar is 50 μm (**b**). Average myotube diameter was measured (n = 3) (**c**). **d**-**f**, Human Skeletal Muscle Myoblasts (HSMM) were differentiated into myotubes and treated with an adenovirus expressing EDA-A2 (**d**) or recombinant EDA-A2 protein (250 ng/ml) (**e**,**f**) for 48 hr. Gene expression was determined by RT-qPCR (n = 3) (**d**,**e**). Human myotubes were investigated under the light microscope. The scale bar is 100 μm (**f**). **g**,**h**, Mouse primary myotubes were treated with recombinant EDA-A1, EDA-A2 or TNFα proteins (250 ng/ml each) for 48 hr. MyHC was immunofluorescently labeled while nuclei was counterstained with DAPI. The scale bar is 50 μm (**g**). MyHC signal was normalized to the number of myotube nuclei (n = 3) (**h**). **i**, Mouse primary myotubes treated with recombinant EDA-A2 (250 ng/ml) and proteasome inhibitor MG132 (10 μM) were lysed and protein samples were studied by western blotting. The values are mean ± SEM. Statistical analysis was conducted using one-way ANOVA (**a**,**c**,**h**) or the two-tailed *t*-test (**d**,**e**). \**p* < 0.05, \*\**p* < 0.01, \*\*\**p* < 0.001, compared with the Ad-LacZ or the Control group.

**Extended Data Figure 4.**
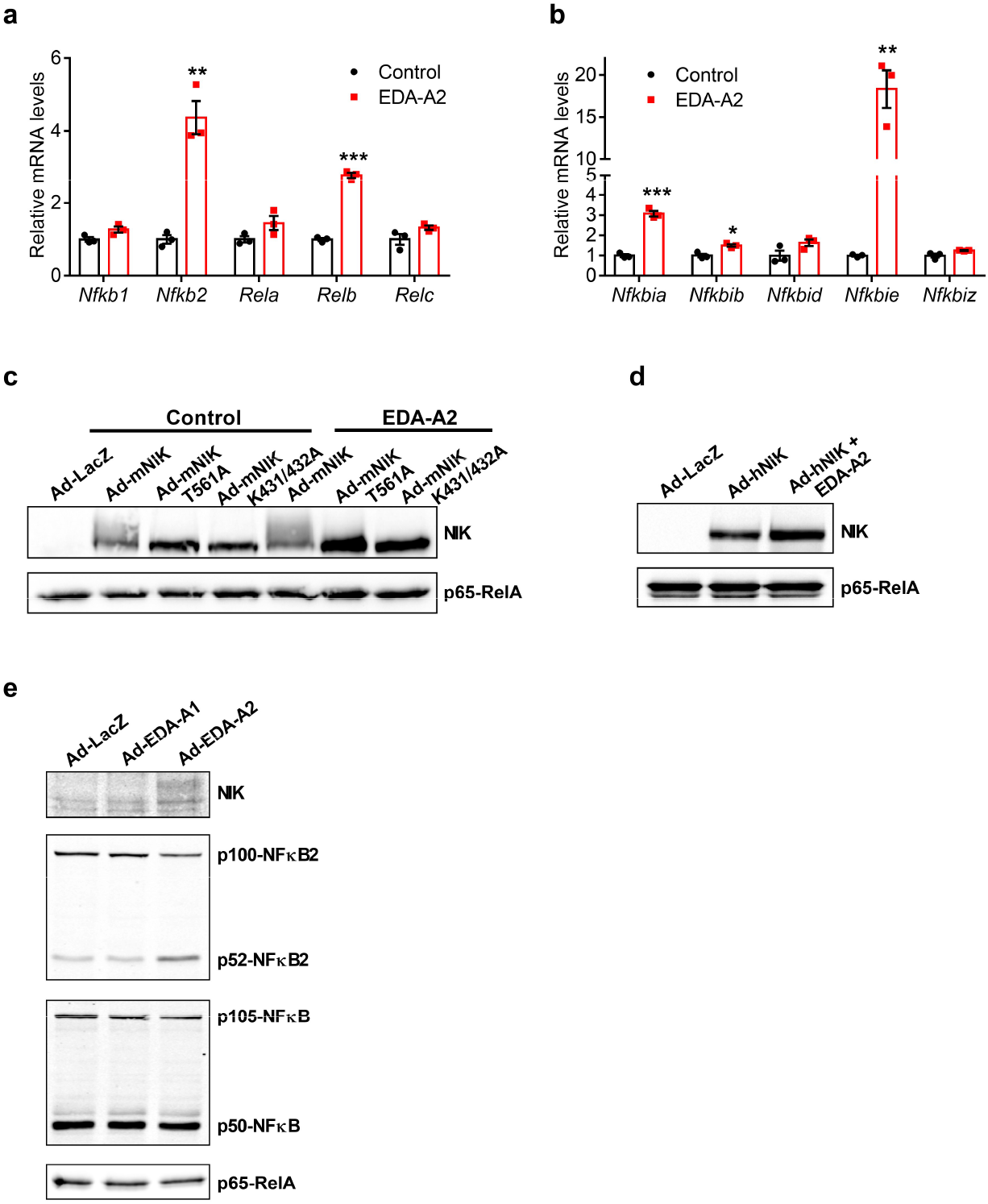
EDA-A2 stimulates the expression of NFκB signaling components and the alternative NFκB activation in primary myotubes. Electrophoretic mobility shift of mouse NIK protein depends on its autophosphorylation and kinase activity. **a**,**b**, Mouse primary myotubes were treated with recombinant EDA-A2 (250 ng/ml) for 24 hr and gene expression was determined by RT-qPCR (n = 3). **c,d**, Mouse primary myotubes were transduced with adenoviruses expressing LacZ, wild-type mouse NIK (mNIK), autophosphorylation-deficient mNIK-T561A mutant and kinase-dead mNIK-K431/432A mutant or human NIK (hNIK). A day later, recombinant EDA-A2 (250 ng/ml) was also added for another 24 hr. Protein levels were determined by western blotting. **e**, Mouse primary myotubes were transduced with LacZ, EDA-A1, or EDA-A2 expressing adenoviruses. 24 hr later, protein levels were determined by western blotting. The values are mean ± SEM. Statistical analysis was conducted using the two-tailed t-test. \**p* < 0.05, \*\**p* < 0.01, \*\*\**p* < 0.001, compared with the Control group.

**Extended Data Figure 5.**
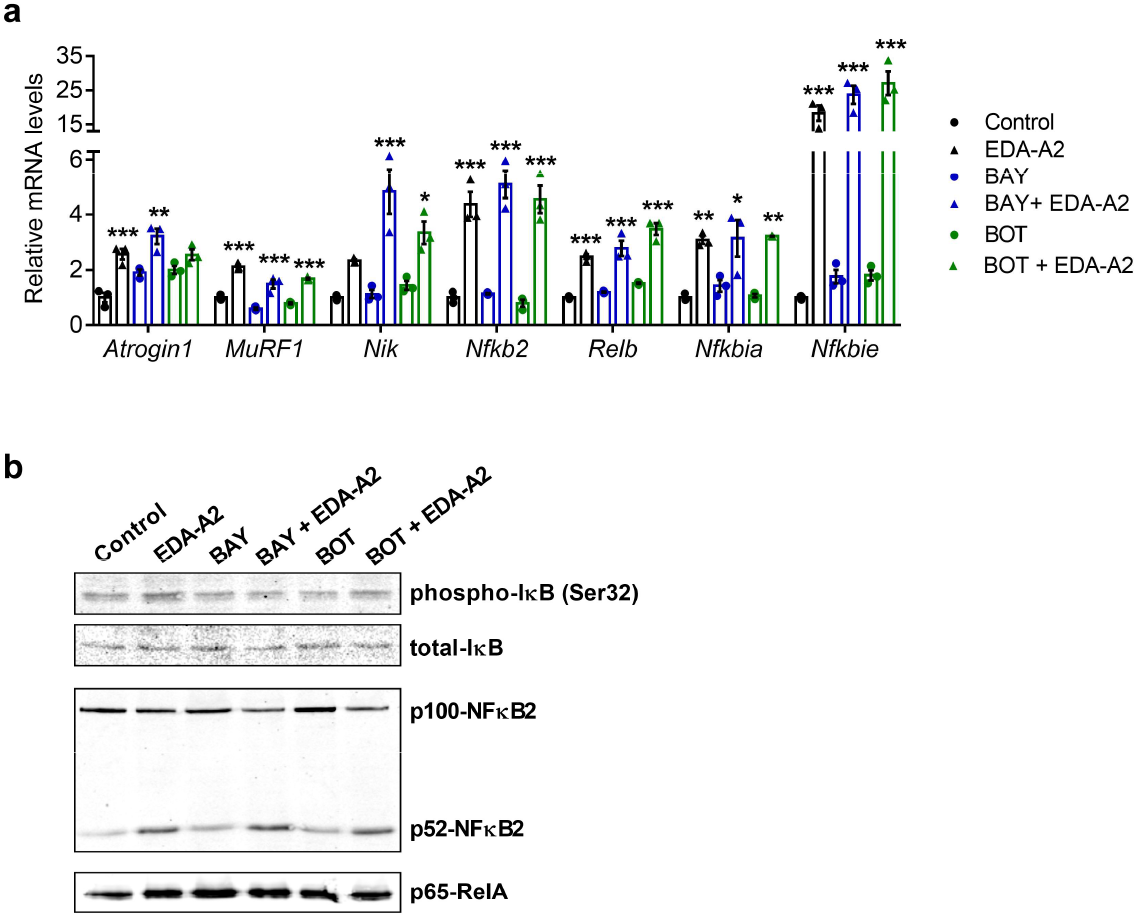
Activation of the canonical NFκB signaling is dispensable for EDA-A2-induced gene expression in primary myotubes. **a**,**b**, Mouse primary myotubes were treated with IκB phosphorylation inhibitors BAY 11-7082 (10 μM) and BOT-64 (10 μM) in combination with recombinant EDA-A2 (250 ng/ml) for 24 hr. Gene expression was studied by RT-qPCR (n = 3) (**a**) and protein levels were determined by western blotting (**b**). The values are mean ± SEM. Statistical analysis was conducted using one-way ANOVA. \**p* < 0.05, \*\**p* < 0.01, \*\*\**p* < 0.001, compared with the control group or the respective inhibitor only group.

**Extended Data Figure 6.**
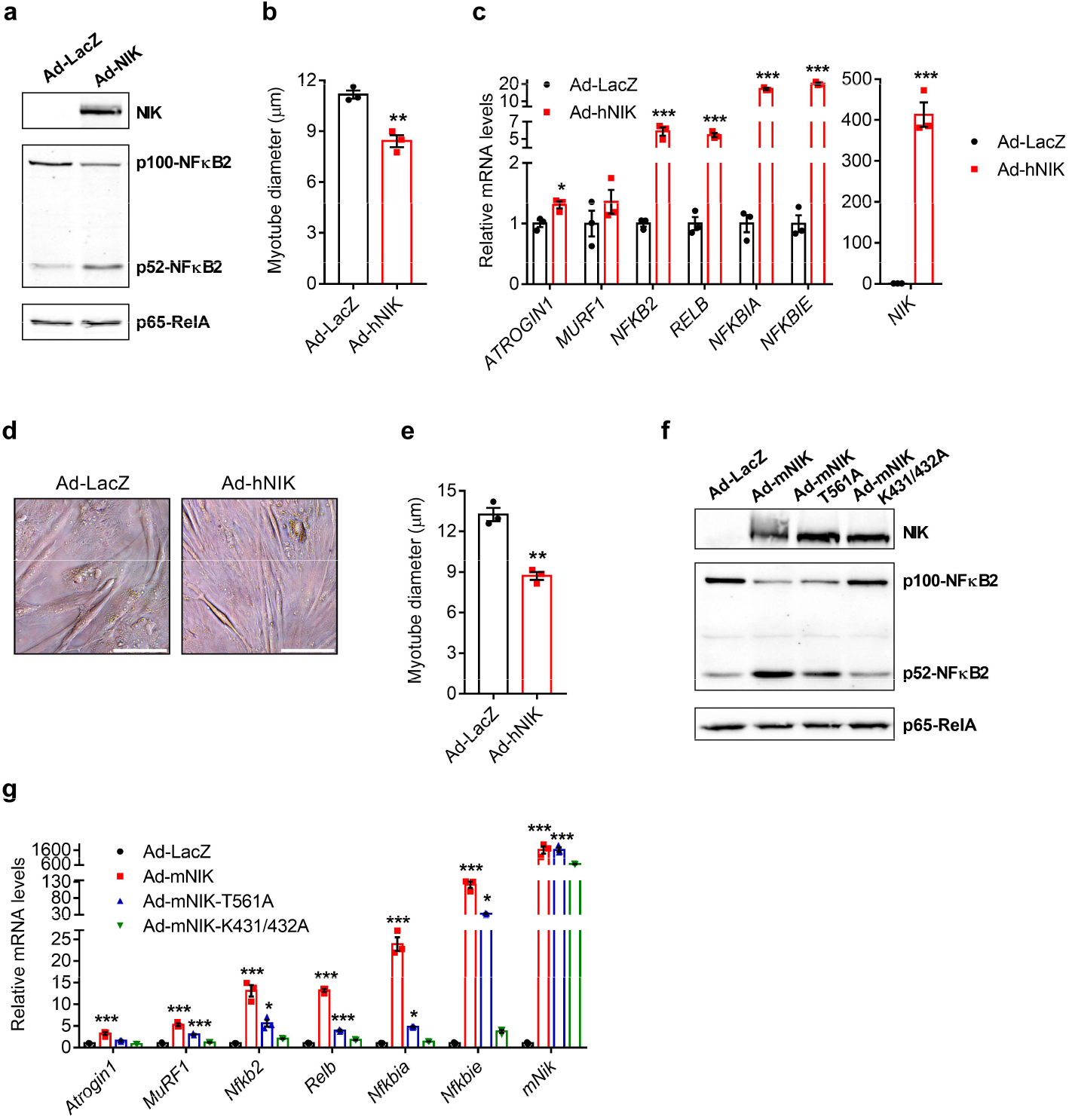
Overexpression of NIK promotes the alternative NFκB activation and atrophy in primary myotubes. **a**,**b**, Mouse primary myotubes were transduced with adenoviruses expressing LacZ or human NIK (hNIK). 24hr later, protein levels were determined by western blotting (**a**). Myotubes also treated with GFP adenovirus were visualized 48 hr later under the fluorescence microscope and average myotube diameter was measured (n = 3) (**b**). **c**-**e**, Human Skeletal Muscle Myoblasts (HSMM) were differentiated into myotubes and treated with adenoviruses expressing LacZ or human hNIK for 48 hr. Gene expression was determined by RT-qPCR (n = 3) (**c**). Human myotubes were investigated under the light microscope. The scale bar is 100 μm (**d**). Myotube diameter was measured (n = 3) (**e**). **f**,**g**, Primary myotubes were transduced with adenoviruses expressing LacZ, mouse NIK (mNIK), autophosphorylation-deficient mNIK-T561A mutant or kinase dead mNIK-K431/432A mutant. Protein levels were determined by western blotting (**f**). This is the same experiment as Extended Data Fig. 4a. NIK and p65-RelA blots were cropped from Extended Data Fig. 4a. mRNA levels were tested by RT-qPCR (**g**). The values are mean ± SEM. Statistical analysis was conducted using the two-tailed t-test (**b**,**c**,**e**) and one-way ANOVA (**g**). \**p* < 0.05, \*\**p* < 0.01, \*\*\**p* < 0.001, compared with the Ad-LacZ group.

**Extended Data Figure 7.**
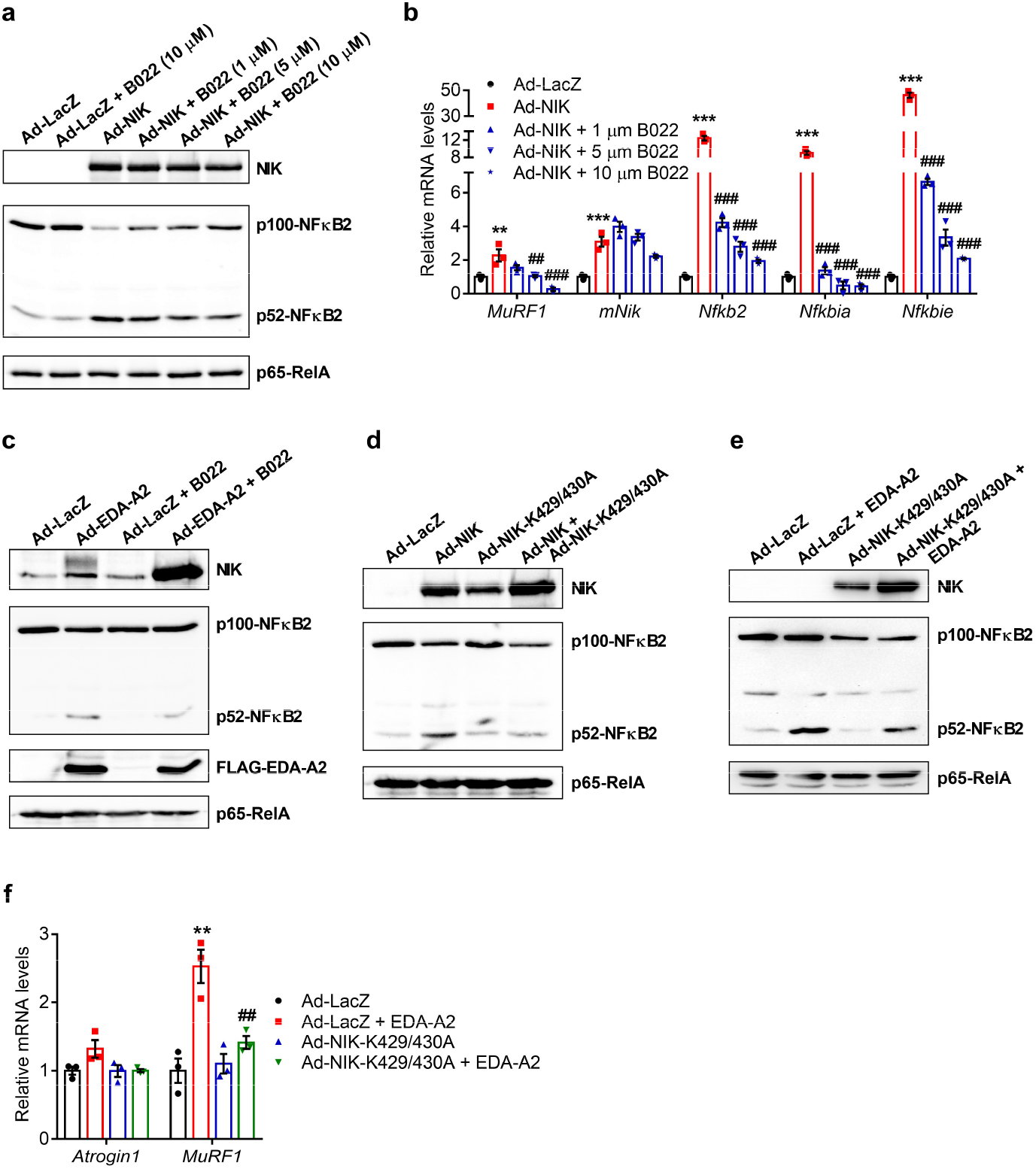
The inhibition of NIK kinase activity with B022 or a dominant-negative NIK mutant blocks EDA-A2’s effects in primary myotubes. **a**,**b**, Mouse primary myotubes were transduced with adenoviruses expressing LacZ or human NIK and treated with different doses of B022 (1 μM, 5 μM or 10 μM) for 24hr. Protein levels were determined by western blotting (**a**) and changes in gene expression were tested by RT-qPCR (n = 3) (**b**). **c**, Mouse primary myotubes were transduced with LacZ or EDA-A2 adenoviruses and treated with B022 (5 μM) for 24hr. Protein levels were determined by western blotting. **d**, Mouse primary myotubes were transduced with adenoviruses expressing wild-type human NIK or the dominant-negative human NIK-K429/430A mutant. Protein levels were determined by western blotting. **e**,**f**, Mouse primary myotubes were transduced with adenoviruses expressing LacZ or the NIK-K429/430A mutant and treated with recombinant EDA-A2 (100 ng/ml) for 24hr. Protein levels were determined by western blotting (**e**) and changes in gene expression were tested by RT-qPCR (n = 3) (**f**). The values are mean ± SEM. Statistical analysis was conducted using one-way ANOVA. \*\**p* < 0.01, \*\*\**p* < 0.001, compared with the Ad-LacZ group. *##p* < 0.01, *###p* < 0.001, compared with the Ad-NIK only group (**b**). *##p* < 0.01 compares differences between EDA-A2 treatment groups (**f**).

**Extended Data Figure 8.**
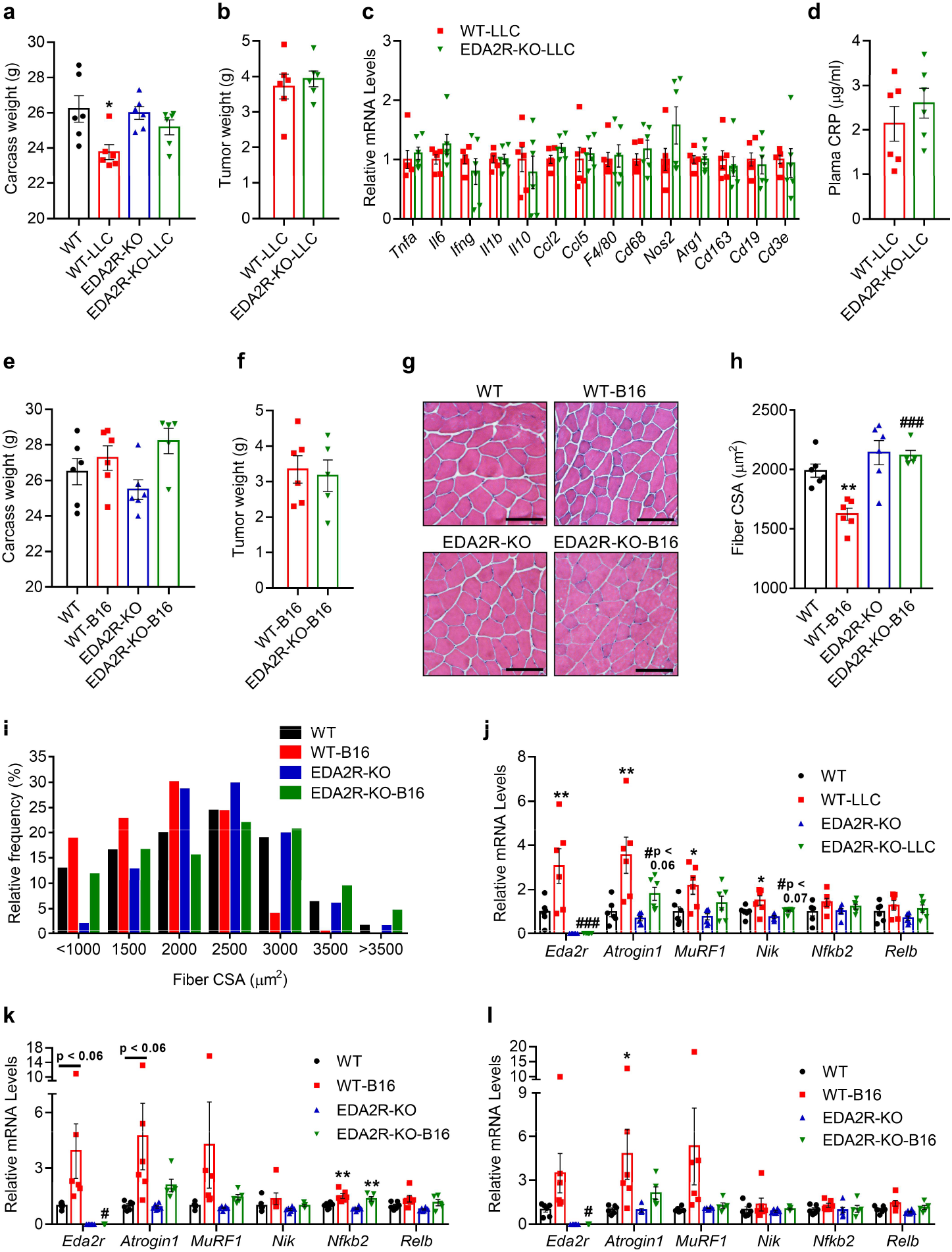
EDA2R-deficient mice are resistant to tumor-driven muscle wasting. **a**-**d**,**j**, Mice were inoculated with LLC cells and sacrificed 16 days later (n = 6). Carcass weight without the tumor mass (**a**) and tumor weight (**b**) were measured. **c**, Gene expression levels in tumor samples were measured by RT-qPCR (n = 6). **d**, Plasma CRP levels were determined by ELISA (n = 6). **e-i**, **k-l**, Mice were inoculated with B16 cells and sacrificed 14 days later (EDA2R-KO-B16 n = 5, other groups n = 6). Carcass weight without the tumor mass (**e**) and tumor weight (**f**) were measured. A decrease in carcass weight was induced by LLC tumors. However, tissue wasting was not reflected in the carcass weight when mice received B16 tumors. Because these tumors cause excessive subcutaneous swelling due to inflammation which masks the wasting. **g-i,**Gastrocnemius muscle cross-sections were H&E stained (**g**), cross-sectional area (**h**) and the fiber frequency distribution (**i**) were measured. The scale bar is 100 μm. **j**, Quadriceps muscle mRNA levels of the LLC tumor-bearing mice were tested by RT-qPCR (n = 6). **k**,**l**, Gastrocnemius muscle (**k**) and quadriceps muscle (**l**) mRNA levels of the B16 tumor-bearing mice were determined by RT-qPCR (EDA2R-KO-B16 n = 5, other groups n = 6). The values are mean ± SEM. Statistical analysis was conducted using two-way ANOVA. \**p* < 0.05, \*\**p* < 0.01 compares differences between tumor-bearing and non-tumor-bearing mice of the same genotype. *#p* < 0.05, *###p* < 0.001 compares differences between tumor-bearing wild-type and tumor-bearing knockout mice.

**Extended Data Figure 9.**
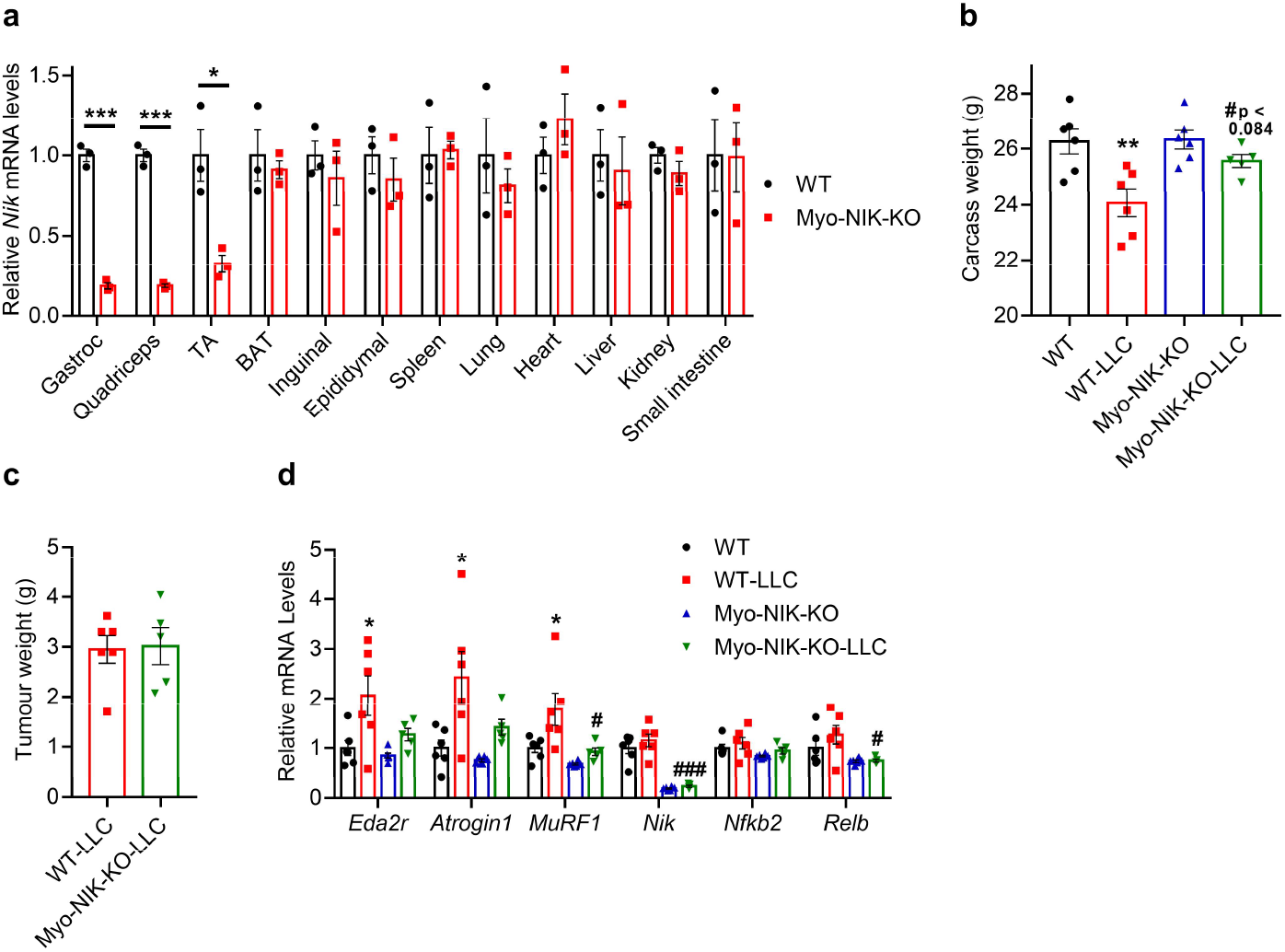
Muscle-specific depletion of NIK protects from tumor-driven muscle wasting. **a**. *Nik*mRNA levels were tested by RT-qPCR in various tissues of the Myo-NIK-KO mice (n = 3). **b-d**, Mice were inoculated with LLC cells and sacrificed 16 days later (Myo-NIK-KO-LLC n = 5, other groups n = 6). Carcass weight without the tumor mass (**b**) and tumor weight (**c**) were measured. Quadriceps muscle mRNA levels were tested by RT-qPCR (Myo-NIK-KO-LLC n = 5, other groups n = 6) (**d**). Statistical analysis was conducted using the two-tailed t-test (**a**) or two-way ANOVA (**b**,**d**). \**p* < 0.05, \*\**p* < 0.01, \*\*\**p* < 0.001 compares differences between WT and WT-LLC groups. *#p* < 0.05, *###p* < 0.001 compares differences between WT-LLC and Myo-NIK-KO-LLC groups.

**Extended Data Figure 10.**
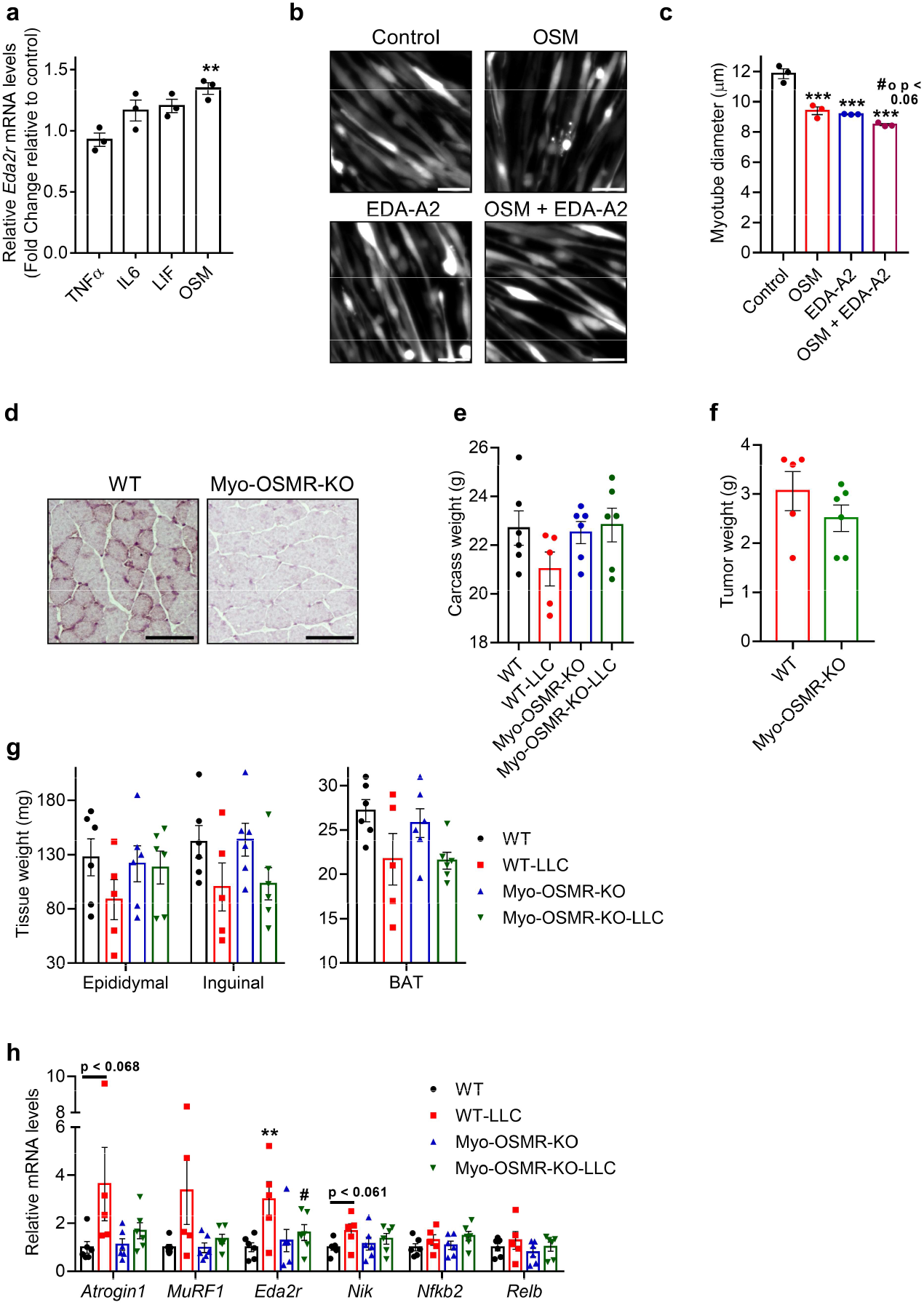
OSM induces *Eda2r* expression in muscle and the depletion of OSMR protects from muscle wasting. **a**, Mouse primary myotubes were treated with recombinant TNFα, IL-6, LIF and OSM (250 ng/ml each). mRNA levels were determined by RT-qPCR (n = 3). **b**-**c**, Mouse primary myotubes were treated with recombinant OSM and EDA-A2 (250 ng/ml each) for 48 hr. Myotubes also treated with the GFP adenovirus were visualized under the fluorescence microscope. The scale bar is 50 μm (**b**). Average myotube diameter was measured (n = 3) (**c**). **d**, OSMR levels in gastrocnemius muscle were determined by immunohistochemistry. **e**-**h**, Mice were inoculated with LLC cells and sacrificed 16 days later (WT-LLC n = 5, other groups n = 6). Carcass weight without the tumor mass (**e**) and tumor weight (**f**) were measured. Collected adipose tissues were weighed (**g**). Quadriceps muscle mRNA levels were tested by RT-qPCR (WT-LLC n = 5, other groups n = 6) (**h**). The values are mean ± SEM. Statistical analysis was conducted using one-way ANOVA (**a**,**c**) and two-way ANOVA (**h**). \*\**p* < 0.01, \*\*\**p* < 0.001 compared to the control group (**a**,**c**). *#op* <0.06 compared to the OSM group (**c**). \*\**p* < 0.01 compares differences between WT and WT-LLC groups *#p* < 0.05 compares differences between WT-LLC and Myo-OSMR-KO-LLC groups (**h**).

